# Decoding molecular and cellular heterogeneity of nucleus accumbens with high-throughput scRNA-seq and MERFISH

**DOI:** 10.1101/2021.07.17.452808

**Authors:** Renchao Chen, Timothy R. Blosser, Mohamed N. Djekidel, Junjie Hao, Aritra Bhattacherjee, Wenqiang Chen, Luis M. Tuesta, Xiaowei Zhuang, Yi Zhang

## Abstract

The nucleus accumbens (NAc) plays an important role in regulating multiple behaviors and its dysfunction has been linked to many neural disorders. However, the molecular, cellular and anatomic heterogeneity underlying its functional diversity remains incompletely understood. Here, we generate a cell census of the mouse NAc using high-throughput single cell RNA sequencing and multiplexed error-robust FISH, revealing a high level of cell heterogeneity in this brain region. We show that the transcriptional and spatial diversity of neuron subtypes underlie NAc’s anatomic and functional heterogeneity, and possibly contribute to the pathogenesis of different neurological disorders. These findings explain how the seemingly simple neuronal composition of the NAc achieves its highly heterogenous structure and diverse functions. Collectively, our study generates a spatially resolved cell taxonomy for understanding the NAc structure and function, which demonstrates the importance of combining molecular and spatial information in revealing the fundamental features of the nervous system.

## Introduction

The nucleus accumbens (NAc) is a key component of the basal ganglion circuitry that plays critical roles in integrating information from cortical and limbic regions to direct different behaviors ^1^. Studies in the past few decades have revealed that the NAc is involved in regulating a wide variety of animal behaviors that include appetitive and aversive responses, feeding, social interaction, as well as reinforcement and instrumental learning ^2–6^. In addition, studies in both human patients and animal models have linked NAc dysfunction to multiple neuropsychiatric disorders, including depression, anxiety, schizophrenia and substance abuse ^7–10^. Consistent with its functional diversity, anatomic analyses revealed that the NAc has complex neural connections with different brain regions. For example, the NAc receives glutamatergic inputs from multiple upstream brain regions, such as the frontal cortex, thalamus, hippocampus, and amygdala, as well as monoaminergic inputs from the ventral tegmental area (VTA), the raphe nucleus and the nucleus of solitary tract. On the other hand, the NAc also projects to different downstream targets, including the ventral pallidum, the lateral hypothalamic area, the VTA, the dorsal raphe and the periaqueductal gray ^11–14^. Accumulating evidence suggests that specific input/output circuits of the NAc may underlie its different functions ^2, 15–18^.

In contrast to its functional diversity and anatomic complexity, our understanding of the cellular composition and architecture of the NAc is still limited. It is generally accepted that over 90% of NAc neurons are GABAergic medium spiny neurons (MSNs), with a small percentage of interneurons (INs) ^19, 20^. The conventional direct/indirect pathway model divides the MSNs into D1 dopamine receptor-expressing and D2 dopamine receptor-expressing subtypes (D1 and D2 MSNs), based on their distinct gene expression, connectivity and function ^20–23^. Although this model provides a framework for understanding the cellular and circuitry organization of the striatum, it takes little consideration of intra-striatal heterogeneity, and thus cannot explain the anatomical and functional diversity of the NAc. For example, the striosome/matrix compartments in the dorsal striatum and the core/shell organization in the NAc have been well recognized, which exhibit discrete molecular and anatomic features ^24, 25^. Additionally, the MSNs located in different or even the same subregions of the NAc are connected with different upstream and downstream targets ^14, 16, 26^. Moreover, different neural functions have been attributed to the NAc subregions, or even the same neuron subtype (e.g. D1 MSN) within close proximity ^16, 27, 28^. In line with this, recent studies have revealed substantial neuronal heterogeneity (especially MSNs) in the striatum with single-cell profiling ^29–31^, but whether molecularly distinct MSN subtypes underlie this anatomical (e.g. core/shell division) and functional heterogeneity of the NAc remains to be shown.

To answer this question, a molecularly defined and spatially resolved cell taxonomy of the NAc is required to integrate its structural and functional complexity. Recent studies have used single cell RNA sequencing (scRNA-seq) and RNA in situ hybridization to characterize the cell composition of the striatum and spatial profiles of marker gene expression, which revealed both discrete and continuous transcriptional programs underlying the MSN heterogeneity, and suggested a correlation between gene expression and spatial distribution of MSNs ^29, 30^. Yet, how cell types contribute to the anatomic and functional heterogeneity of the NAc remains an open question, and a systematic characterization of the spatial distributions of cell types could help provide an answer. In this study, we analyzed the mouse NAc at single-cell resolution by combining scRNA-seq and multiplexed error-robust FISH (MERFISH) ^32, 33^, which not only revealed molecularly distinct neuron subtypes, but also resolved how these cell types are spatially organized into NAc subregions. Furthermore, integrative analyses of the molecular and spatial features of NAc neuron subtypes enabled us to link different functions to different neuron substrates of the NAc, illuminating the underlying molecular-cellular-anatomical-functional relationship of the NAc. Collectively, our study generated a molecular and anatomic taxonomy of the NAc, which could serve as a basis for understanding how different neuron subtypes of this brain region are involved in various physiological and pathological conditions.

## Results

### Single-cell RNA-seq reveals the major cell populations in the NAc

To systematically characterize the cell types of the mouse NAc, we performed high-throughput single-cell RNA-seq with cells dissociated from the NAc of adult mice (**Fig. 1a** **and Extended Data Fig. 1a**). In total, eleven independent biological samples were collected (**Extended Data Fig. 1b and Supplementary Table 1**), with a median of 5,084 single cells captured in each sample. After filtering out low-quality cells (< 1,500 gene detected/cell) and cells with high mitochondrial reads (> 10%), we obtained 37,011 single cells. Our initial “low-resolution” clustering analysis revealed nine major cell clusters with distinct gene expression patterns, including four neuronal populations (*Snap25^+^*) and five non-neuronal populations (*Snap25*^-^) (**Fig. 1b-d** **and Supplementary Table 2**), which are present in all eleven samples (**Extended Data Fig. 1c**). The five non-neuronal populations identified from the NAc are highly similar to those identified from other brain regions ^34–36^, including astrocytes, immune cells, blood vessel cells, oligodendrocyte progenitor cells (OPC) and oligodendrocytes (**Fig. 1b-d**), which are consistent with their broad distribution in the brain. Since the numbers of UMIs (unique molecular identifier) and genes detected in non-neuronal cells are significantly lower than those detected in neuronal cells (**Extended Data Fig. 1d**), we further included non-neuronal cells with 800-1500 genes detected in each cell by building a predictive model to assign these cells to corresponding cell clusters (**Extended Data Fig. 1e, f**), which increased the cell number to 47,576, with a median UMI of 4,867 and a median of 2,067 detected genes in each cell. For neuronal cells, we identified D1 and D2 MSNs as the major neuron populations in the NAc, which constitute more than 90% of neurons (20,029 out of 21,842 neurons), as well as a relatively small interneuron (IN) population (**Fig. 1b-d**). Interestingly, one cell population representing neural stem cell and neuroblast was identified (**Fig. b-d**) in the ventral wall of the lateral ventricle of the adult mouse brain (**Extended Data Fig. 1g**). To characterize the neuronal heterogeneity of the NAc, we focused our further analysis on IN and MSN, respectively.

**Fig. 1.**
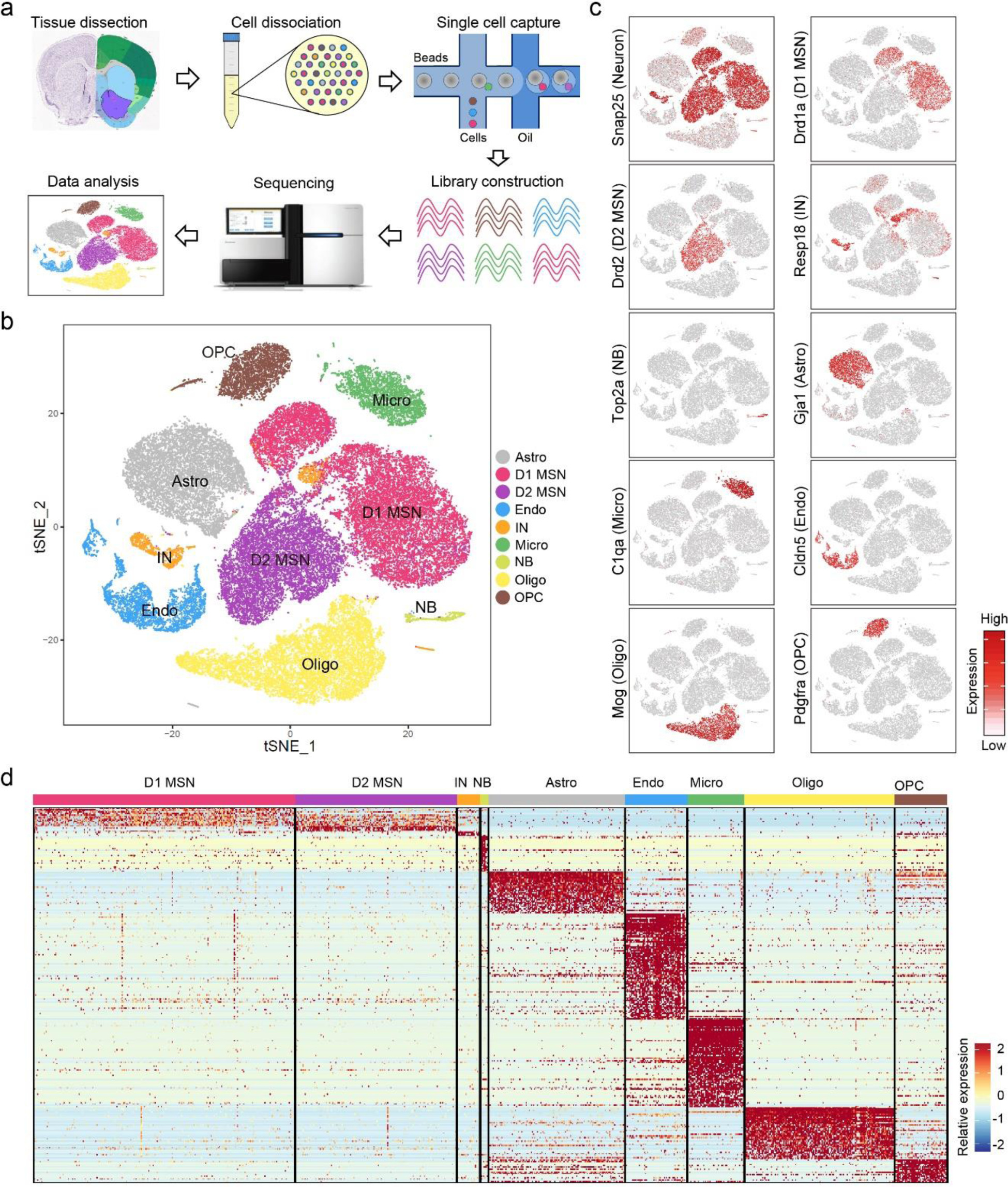
Single-cell RNA-seq reveals major cell populations in NAc. **a,** Workflow of single-cell RNA-seq of mouse nucleus accumbens. NAc tissues were dissected from adult mouse brain and dissociated into single-cell suspension. Single cells were captured into droplets with the 10X platform, followed by cDNA synthesis, amplification and library construction. After sequencing, cells were classified by their transcriptomes. **b,** tSNE plot showing the different major cell types in NAc. Different cell clusters are color coded. Astro, astrocyte; D1 MSN, D1-type medium spiny neuron; D2 MSN, D2-type medium spiny neuron; Endo, endothelial cell; IN, interneuron; Micro, microglia; NB, neural stem cells and neuroblast; Oligo, oligodendrocyte; OPC, oligodendrocyte progenitor cell. **c,** tSNE plots showing expression of cell type-specific markers across different cell subtypes. The gene expression level is color-coded. **d,** Heatmap showing the cell type-specific markers are differentially expressed across the 9 NAc cell populations. Differentially expressed genes with power > 0.4, fold change > 2 among the 9 cell clusters were used to generate the heatmap. Columns represent individual cells and rows represent individual genes. The gene expression level is color-coded.

### Transcriptionally defined IN subtypes and their distribution in the NAc

Although INs represent <10% of the total neuron population in the striatum, they exhibit diverse morphological and electrophysiological properties ^19^ and play important roles in regulating the local circuitry ^37, 38^. However, a detailed molecular classification of the NAc IN has not yet been achieved. Further clustering of the initial IN population identified 13 subtypes with distinct transcriptional features (**Fig. 2a, b** **and Extended Data Fig. 2a, b**). The marker genes detected by scRNA-seq for different IN subtypes could be confirmed by ISH (**Extended Data Fig. 2c**). Examination of the expression of conventional striatal IN markers (*Chat, Pvalb, Sst, Th*) in our IN clusters revealed that some of these markers labeled single IN subtypes, such as *Chat* and *Pvalb* (**Fig. 2b** **and Extended Data Fig. 2b, d**). However, scRNA-seq also allowed us to further classify conventional IN populations into different subtypes. For example, we found that the *Sst*^+^ INs could be further divided into *Hhip*^+^ and *Hhip*^-^ subtypes (**Fig. 2b** **and Extended Data Fig. 2b, d**), while the *Th*^+^ INs can be further classified into *Trh*^+^ and *Calb2*^+^ subtypes (**Fig. 2b** **and Extended Data Fig. 2b, d**). These IN subtypes revealed by scRNA-seq may underlie the morphological and electrophysiological heterogeneity observed in Sst^+^ and Th^+^ INs ^19^. Interestingly, we found *Pthlh*, a newly identified marker for cortical chandelier cells ^39^, is also a pan-marker for three NAc IN subtypes exhibiting similar transcriptomes (*Pthlh*^+^/*Pvalb*^+^, *Pthlh*^+^/*Crhbp*^+^ and *Pthlh*^+^/*Pvalb*^-^/*Crhbp*^-^ subtypes) (**Fig. 2b** **and Extended Data Fig. 2b, d**). On the other hand, *Afap1* marks an IN subtype expressing *Npy* but not *Sst* (cluster 8) (**Fig. 2b** **and Extended Data Fig. 2b**), which likely represents the *Npy*^+^ neurogliaform cells ^40^. The IN subtypes identified here are highly similar to those reported in a recent study focusing on IN populations in the dorsal striatum ^41^.

**Fig. 2.**
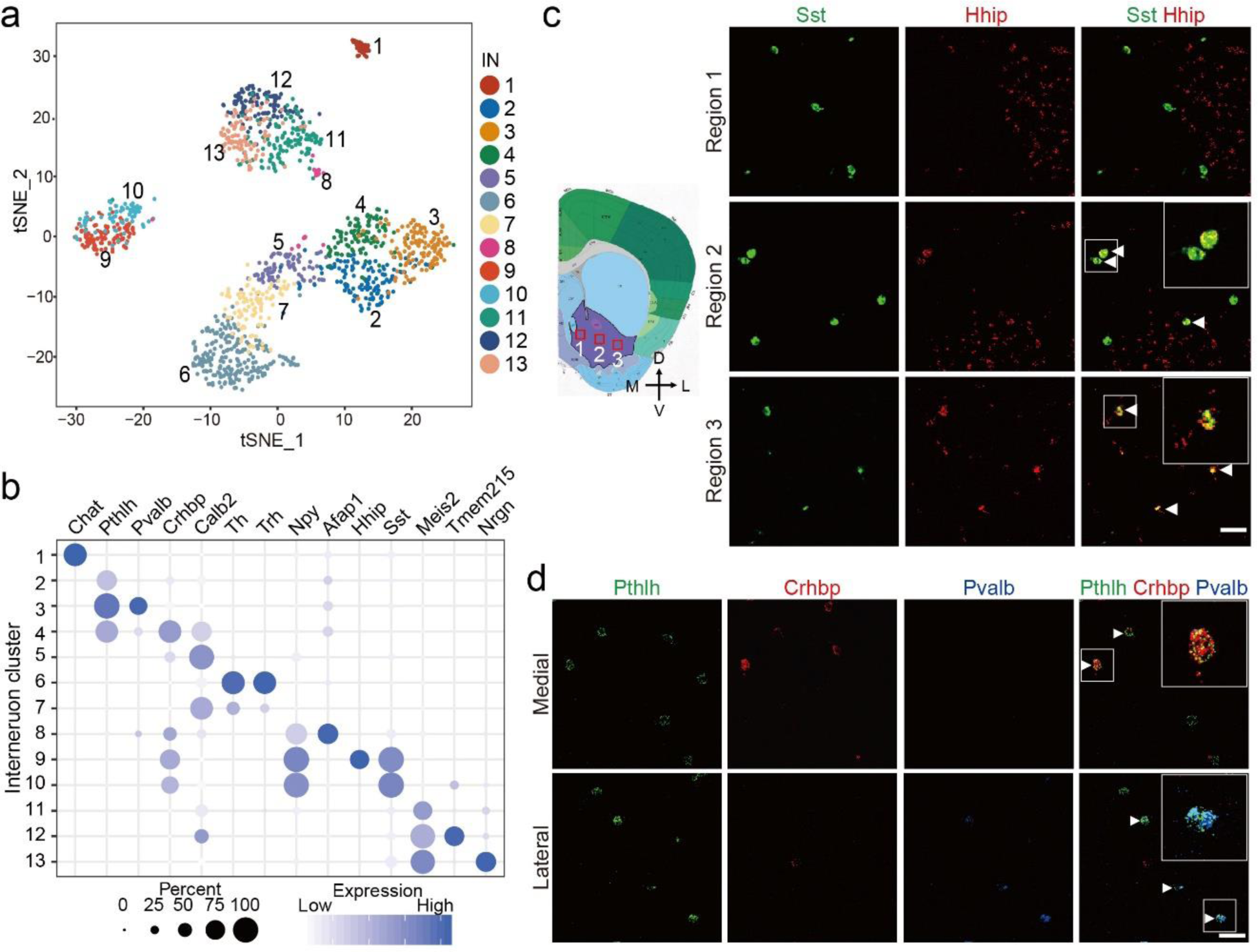
Gene expression and spatial pattern of NAc interneuron subtypes. **a,** tSNE plot showing the 13 interneuron subtypes identified in NAc. Differentially expressed genes among all subtypes are used for dimension reduction. Different neuron subtypes are color coded. **b,** Dot plot showing the expression of selective markers in different NAc interneuron subtypes. The diameters of the dots represent the percentage of cells within a cluster expressing that gene. The gene expression level is color-coded. **c,** FISH showing the distribution of *Sst*^+^/*Hhip*^+^ and *Sst*^+^/*Hhip*^-^ interneuron subtypes in different sub-regions of NAc. The box regions labeled with 1 to 3 in the left panel indicate different sub regions of NAc (from medial to lateral), which are analyzed in the right panels (from upper to lower). The right panels showing the FISH of NAc slice with *Sst* and *Hhip* probes. Arrow heads indicated cells co-expressing *Sst* and *Hhip*. Scale bar100 µm. **d,** FISH showing the enrichment of distinct *Pthlh*^+^ interneuron subpopulations in different sub regions of NAc. The upper and lower panels represent the medial and lateral regions of NAc. Triple-color FISH was performed with probes targeting *Pthlh*, *Crhbp* and *Pvalb*. Arrow heads indicate cells co-expressing *Pthlh* and *Crbhp* (upper panels) or cells co-expressing *Pthlh* and *Pvalb* (lower panels). Scale bar, 50 µm.

By performing multi-colored fluorescent *in situ* hybridization (FISH), we confirmed the identity of *Th*^+^/*Calb2*^+^, *Th*^+^/*Trh*^+^, *Sst*^+^/*Hhip*^+^, *Sst*^+^/*Hhip*^-^, *Pthlh*^+^/*Pvalb*^+^, *Pthlh*^+^/*Crhbp*^+^ and *Pthlh*^+^/*Pvalb*^-^ /*Crhbp*^-^ IN subtypes in the NAc (**Fig. 2c, d** **and Extended Data Fig. 2e**). Notably, different IN subtypes exhibit unique sub-region-specific distribution in the NAc. For example, *Sst*^+^ INs are distributed throughout the NAc (**Fig. 2c**), while the *Sst*^+^/*Hhip*^-^ and *Sst*^+^/*Hhip*^+^ IN subtypes are preferentially located in the medial and lateral NAc, respectively (**Fig. 2c**). Similarly, one *Pthlh*^+^ IN subtype, the *Pthlh*^+^/*Pvalb*^+^ INs, is located mainly in the lateral NAc (**Fig. 2d**), while the other subtype, *Pthlh*^+^/*Crhbp*^+^ neurons, is mainly located in the medial NAc (**Fig. 2d**). Collectively, our scRNA-seq and FISH analysis revealed that NAc has a rich IN diversity with distinct transcriptional and spatial features.

### Transcriptionally defined MSN subtypes exhibit distinct spatial patterns in the NAc

Based on the expression of canonical MSN marker *Ppp1r1b* ^42^, most neuronal cells captured in our scRNA-seq experiments belong to MSNs (**Fig. 3a**). Consistent with the classic direct/indirect pathway model ^20^, the entire MSN population can be divided into discrete D1 and D2 groups based on the expression of known markers, such as *Drd1*, *Pdyn*, *Drd2* and *Adora2a* (**Fig. 3a**). To further understand the heterogeneity within the D1 and D2 MSNs, we separately classified the D1 and D2 MSN populations into 8 subtypes each (**Fig. 3b** **and Extended Data Fig. 3a**).

**Fig. 3.**
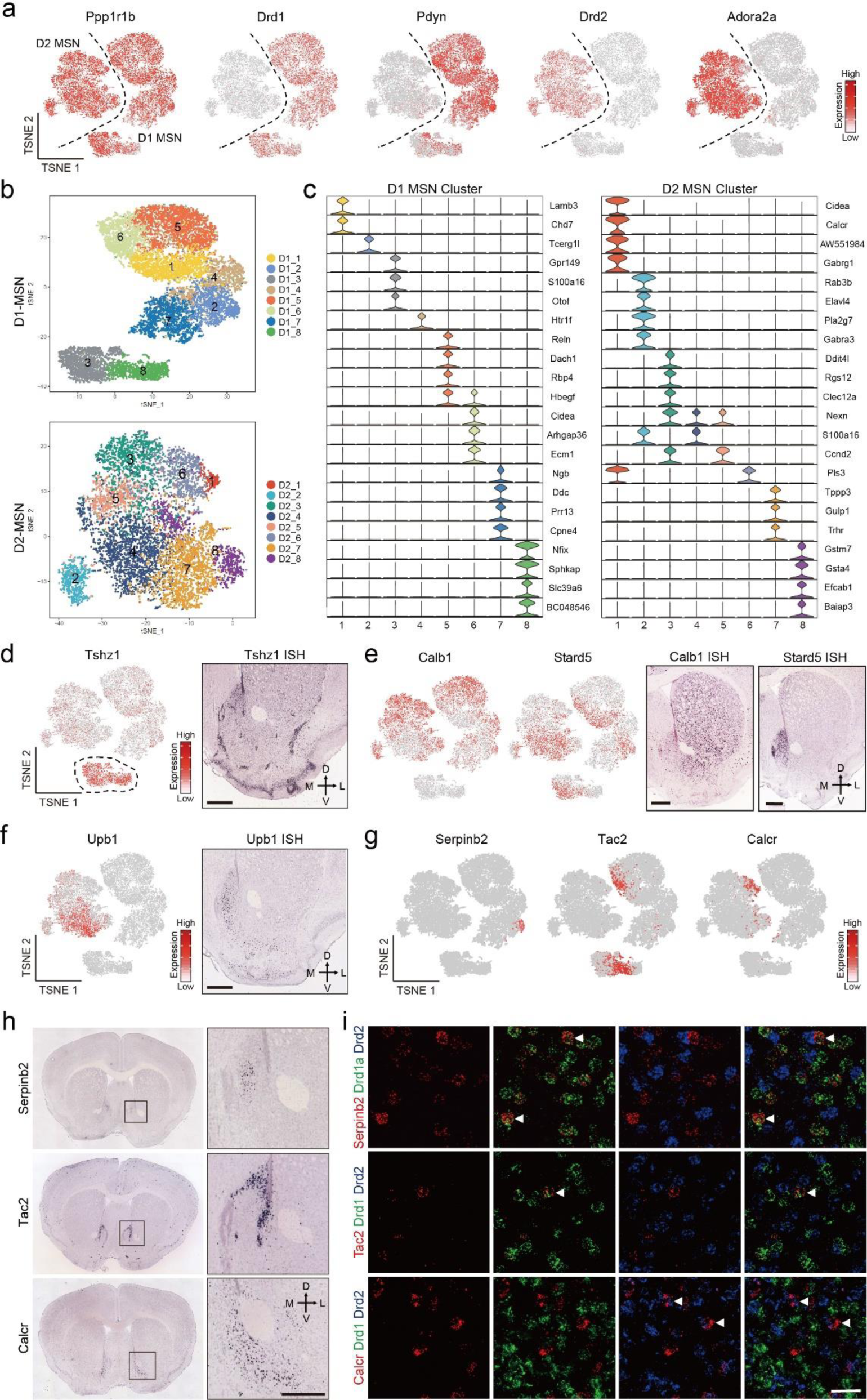
Transcriptional features of MSN subtypes correlate with their spatial distribution in NAc. **a,** tSNE plots showing the expression pattern of *Ppp1r1b*, *Drd1*, *Pdyn*, *Drd2* and *Adora2a* in different MSN populations. The expression level is color-coded. **b,** tSNE plots showing the eight D1 MSN subtypes (upper panel) and eight D2 MSN subtypes (lower panel), different subtypes are color-coded. **c,** Violin plots showing the expression pattern of MSN subtype-specific gene markers across the D1 MSNs (left panel) and D2 MSNs (right panel). Different MSN subtypes are color-coded. The mRNA level is presented on a log scale and adjusted for different genes. **d,** *Tshz1* is enriched in a subpopulation of D1 MSNs. Left panel, tSNE plots showing the expression pattern of *Tshz1* across MSN populations. The expression level is color-coded. Right panel, in situ hybridization of *Tshz1* showing the distribution of *Tshz1*^+^ cells in NAc. Coronal section of mouse brain including NAc is shown. Data is obtained from Allen Mouse Brain Atlas. Scale bar, 500 µm. **e,** Calb1 and Stard5 showing anti-correlated expression pattern. Left two panels, tSNE plots showing the expression of Calb1 and Stard5 as detected by scRNA-seq across D1 and D2 MSNs. The gene expression level is color-coded. The right two panels, *in situ* hybridization of *Calb1* and *Stard5* showing their anti-correlated expression pattern in NAc. The ISH data are from Allen Brain Atlas. Scale bars, 500 µm. **f,** *Upb1*^+^ D2 MSNs are distributed in the medial NAc. Left panel, tSNE plot showing *Upb1* expression is restricted in certain D2 MSN population. Right panel, ISH image showing the distribution of *Upb1*^+^ cells in NAc. Data is obtained from Allen Mouse Brain Atlas. Scale bar, 500 µm. **g,** tSNE plots showing the expression of selected MSN subtype markers across MSNs. Transcriptional levels are color-coded. **h,** ISH images showing the spatial distribution of *Serpinb2*, *Tac2* and *Calcr,* markers of certain D1/D2 MSN subtypes, in mouse NAc. Boxed regions in left panels are enlarged and shown in the right panels. The data were from the Alen Mouse Brain Atlas. Scale bars, 500 µm. **i,** Three-color FISH confirms the expression of MSN subtype-specific markers in NAc. The arrowheads indicate cells that co-express selected MSN subtype markers with *Drd1* or *Drd2* in mouse NAc. Scale bar, 50 µm.

Consistent with a previous report ^30^, most D1 and D2 MSN subtypes show continuous transcriptional features (**Fig. 3b**), as most of the adjacent clusters within D1 and D2 were not separated by clear gaps. Although different subtypes within D1 or D2 could be distinguished by the enriched expression of a panel of marker genes (**Fig. 3c**), these subtypes could represent populations of cells with a continuous spectrum.

Analysis of the *in situ* expression of marker genes revealed that different MSN subtypes tend to locate in different NAc subregions. For instance, we found a subgroup of D1 MSNs (Cluster 3 and 8) express high levels of *Tshz1*, *Lrpprc* and *Foxp2*, but low levels of *Foxp1* and *Isl1* (**Fig. 3d** **and Extended Data Fig. 3b**). *In situ* hybridization (ISH) indicated that the *Tshz1*-high cells are mainly distributed along the boundary of NAc and formed clusters within the NAc (**Fig. 3d** **and Extended Data Fig. 3c**). On the other hand, *Ddit4l*, which labels D1 cluster 4, D1 cluster 5 and D2 cluster 3, is selectively expressed in the dorsolateral part of the NAc (**Extended Data Fig. 3d**), while *Trhr*, which mainly marks the D1 cluster 1, D1 cluster 6 and D2 cluster 7, is selectively expressed in the medial part of the NAc (**Extended Data Fig. 3e**).

To assess the general relationship between transcriptional feature and spatial distribution of MSN subtypes, we grouped the highly variable genes among MSN clusters based on their co expression (**Extended Data Fig. 3f**) and then checked the expression pattern of different gene groups in Allen Brain Atlas. We found that different groups of variable genes exhibit different spatial patterns in the NAc, suggesting that the transcriptional profiles of MSN subtypes are correlated with their anatomical distribution in the NAc. Notably, many spatially related genes are broadly expressed in both D1 and D2 MSNs, indicating that similar transcriptional program underlies the distribution patterns in both D1 and D2 MSNs (such as dorsal-ventral, medial lateral pattern) ^30^. For example, *Calb1* and *Stard5* belong to two anti-correlated gene groups, and each of them is expressed in multiple D1 and D2 MSN subtypes (**Fig. 3e**). ISH analysis demonstrated a clear bias of *Calb1* in lateral, and *Stard5* in the medial NAc (**Fig. 3e**) ^30^. On the other hand, there are also spatially related genes that are restricted to either D1 or D2 subtypes, which may underlie the spatial pattern of certain D1 or D2 MSN subtypes. For example, we found *Upb1* is selectively expressed in a few D2 MSN subtypes located in the medial NAc (**Fig. 3f**).

Taking advantage of the large number of MSNs sequenced in our dataset, we further classify the D1 and D2 MSNs into 30 D1 and 27 D2 subtypes. These “fine” D1 and D2 MSN subtypes could be distinguished in a few instances by unique markers but otherwise mostly by a combination of markers (**Extended Data Fig. 3g, h**). We analyzed the expression of some genes that uniquely mark MSN subtypes, and found that these “fine” MSN subtypes also exhibit distinct spatial patterns, even in restricted NAc subregions. For example, *Serpineb2*, which selectively marks D1 MSN subtype 24 (**Fig. 3g** **and Extended Data Fig. 3g**), is specifically expressed in a small cell population in the medial NAc (**Fig. 3h**) ^30^. Similarly, another D1 MSN subtype (subtype 10), marked by *Tac2* (**Fig. 3g** **and Extended Data Fig. 3g**), is also restricted to the medial NAc (**Fig. 3h**). However, the spatial patterns of these two subtypes are clearly different (**Fig. 3h**). On the other hand, *Calcr*, a marker for D2 MSN subtype 21 (**Fig. 3g** **and Extended Data Fig. 3h**), is expressed selectively in cells located around the anterior commissure (**Fig. 3h**). Using three color FISH, we confirmed that *Serpinb2* and *Tac2* are selectively expressed in the *Drd1*^+^ neurons, while *Calcr* is only expressed in the *Drd2*^+^ neurons (**Fig. 3i**), confirming that they belong to distinct MSN subtypes. Collectively, these results not only reveal a tremendous MSN heterogeneity within the NAc, but also suggest a relationship between gene expression and spatial features of certain neuron subtypes, which may underlie the anatomical and functional heterogeneity of NAc subregions ^6^.

### Mapping transcriptionally distinct cell types of the striatum with MERFISH

The scRNA-seq and ISH analyses described above indicate a correlation between transcriptional and spatial features of different neuron subtypes in the NAc. However, the technical limitation of conventional ISH prevented us from carrying out a comprehensive spatial distribution analysis of all the cell subtypes, which requires analysis of sufficient genes at single-cell resolution in brain slices. To this end, we performed MERFISH, a single-cell transcriptome imaging method^32, 33^, to systematically map the spatial patterns of different cell subtypes across the NAc.

For MERFISH experiments, we selected a panel of 253 gene targets (**Supplementary Table 3**), including marker genes for major cell populations and neuron subtypes identified by scRNA-seq, as well as genes with strong functional implications in the NAc (such as channels, neuropeptides and receptors). Among the 253 genes, 251 were detected by combinatorial error-robust barcodes with MERFISH imaging, while the remaining 2 genes were detected with sequential imaging rounds due to their high expression levels (**Fig. 4a**). To cover the NAc region along the anterior posterior (AP) axis, we performed MERFISH imaging on serial coronal sections collected between Bregma 1.94 mm and 0.74 mm (with 100um interval between consecutive slices) of the adult mouse brain (10-week old) (**Fig. 4a** **and Extended Data Fig. 4a**). The imaging area spans the entire striatum as no clear anatomic features could be used to accurately separate the NAc from other striatal divisions.

**Fig. 4.**
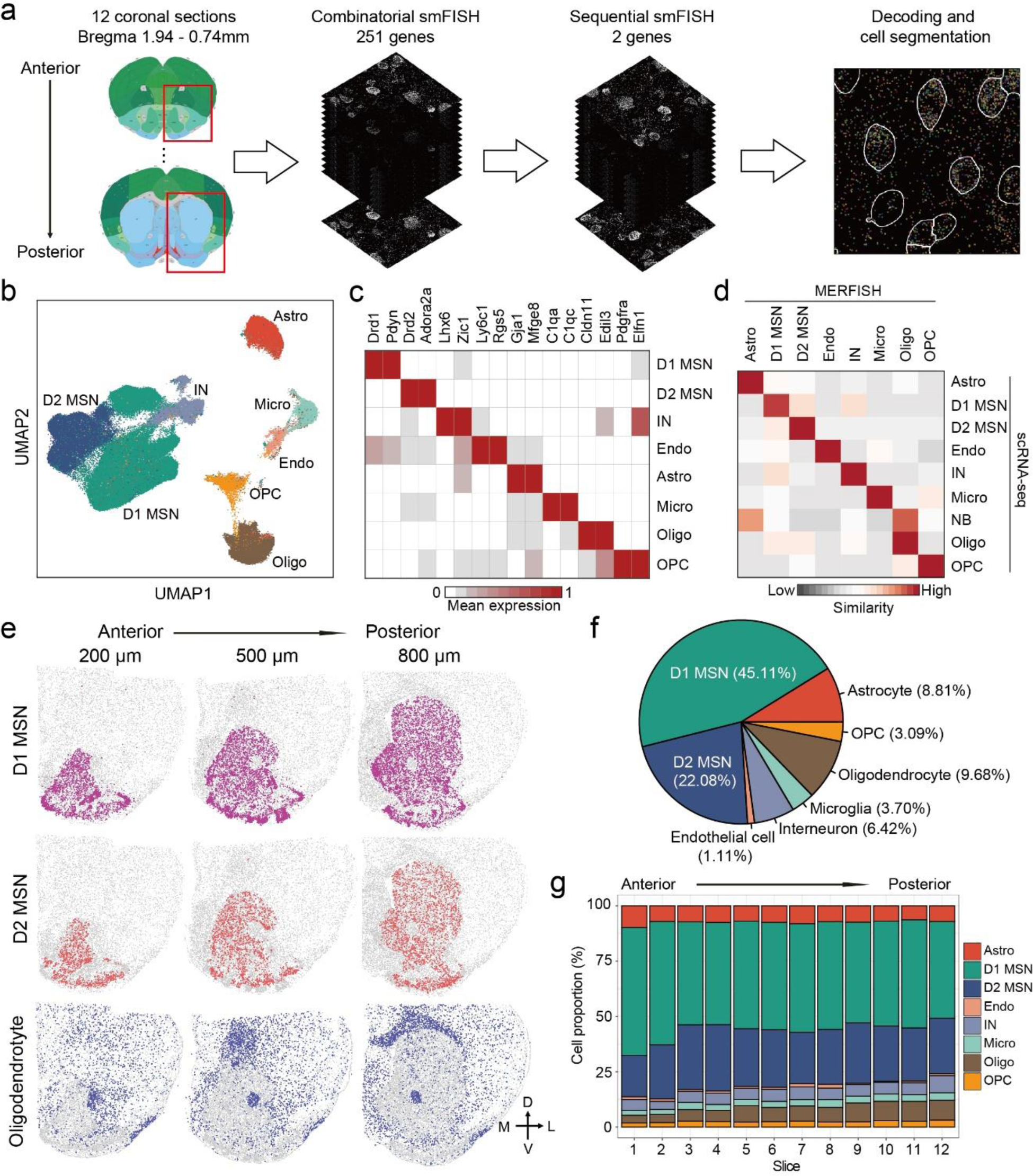
Mapping transcriptionally distinct cell types in mouse striatum with MERFISH. **a,** The workflow of MERFISH profiling of striatum tissue. **b,** UMAP plot showing the major striatal cell populations identified by MERFISH. The different cell populations are color coded. **c,** Heatmap showing the expression pattern of selected marker genes across the major striatal cell populations. The expression level is color-coded. **d,** Heatmap showing the correspondence between scRNA-seq and MERFISH on classification of major striatal cell types. The similarity between scRNA-seq and MERFISH cell clusters is defined as the proportion scRNA-seq cells that could be matched to each MERFISH cluster for each scRNA-seq cluster (see Methods for details). **e,** Spatial distribution of D1 MSN, D2 MSN and oligodendrocyte in brain slices at different anterior-posterior positions. Three of the twelve slices from a male mouse were shown. Colored dots were cells belong to the specified cell populations, while gray dots indicate all other cells. The 200, 500 and 800 μm labels indicate the distance from the anterior position (Bregma 1.94mm). The dorsal-ventral (DV) and medial-lateral (ML) axes were illustrated. **f,** Pie chart showing the percentages of the major striatal cell populations detected by MERFISH. g, Bar graph showing the percentage of the major cell populations across the twelve brain slices along the anterior-posterior (AP) axis.

Individual mRNA molecules were readily detected, decoded to determine their gene identities and assigned to individual cells segmented based on DAPI and Poly A staining (**Extended Data Fig. 4b, c**), thus generating the expression profile of target genes across all analyzed cells (**Fig. 4a**). Of the two replicates, MERFISH showed high consistency (r=0.96) in detecting the mean number of different mRNA molecules (**Extended Data Fig. 4d**), and the gene expression level determined by MERFISH correlated well with that of bulk RNA-seq (**Extended Data Fig. 4e**). In addition, the spatial patterns of certain genes determined by MERFISH are consistent with that revealed by conventional ISH (**Extended Data Fig. 4f**), validating the reliability of the analyses.

In total, we profiled over 700,000 cells in two biological replicates. Clustering analysis with the single-cell gene expression profile obtained by MERFISH identified major cell populations representing D1 and D2 MSN, IN, astrocyte, endothelial cell, OPC, oligodendrocyte and microglia (**Methods and** **Fig. 4b**) based on the expression of known marker genes (**Fig. 4c**). These major cell populations and their relative abundance exhibited little bias between the two MERFISH replicates (**Methods and Extended Data Fig. 4g, h**), suggesting no significant batch effect. In addition, the cell types revealed by MERFISH are consistent with those determined by scRNA-seq (**Methods and** **Fig. 4d**), suggesting that MERFISH reliably captured molecularly distinct cell populations. Importantly, the spatial patterns of major cell types revealed from MERFISH are well in line with known anatomic features of this brain regions: 1) The D1 and D2 MSNs span the entire striatal structure, including the dorsal striatum, the NAc and the olfactory tubercle (OT) (**Fig. 4e**); 2) The oligodendrocytes are highly enriched in the anterior commissure (AC) and corpus callosum, with others dispersed across the striatum (**Fig. 4e**); 3) INs, astrocytes, OPCs, endothelial cells and microglia are dispersed across the entire striatum (**Extended Data Fig. 4i**). These results demonstrated the capacity of MERFISH in resolving spatial patterns of molecularly defined cells *in situ*. Based on the MERFISH result, we estimated that striatum contains about 45% D1 MSN, 22% D2 MSN, 6% interneuron and 27% non-neuronal cells (**Fig. 4f**). The relative abundance of different neuron and non-neuron populations is largely stable across different coronal sections (**Fig. 4g**), suggesting that the major cell populations in the striatum are evenly distributed along the AP axis.

### MSN subtypes revealed by MERFISH are largely consistent with that from scRNA-seq

Our scRNA-seq analysis revealed substantial neuronal heterogeneity in the NAc, but how these molecularly define neuron subtypes contribute to its anatomic and functional properties is largely unknown. To reveal the spatial distribution patterns of different neuronal subtypes, we further classify the D1 and D2 MSN population into 15 D1- and 9 D2-subtypes based on the MERFISH dataset (**Fig. 5a, b**). Similar to scRNA-seq, we found many subtypes within D1 or D2 form a continuous transcriptional spectrum (**Fig. 5a, b**), consistent with a previous report ^30^.These subtypes were named based on their spatial patterns (DS, dorsal striatum; NA, nucleus accumbens; OT, olfactory tubercle; AT, atypical). The similar UMAP distribution and relatively stable ratio for each of the D1 and D2 subtypes between the two MERFISH replicates suggest no significant batch effect (**Extended Data Fig. S5a-c**). Different D1 and D2 MSN subtypes showed enriched expression of different gene sets, but the majority of them do not have a single distinguishable marker (**Fig. 5c, d**, **Extended Data Fig. 5d, e and Supplementary Table 4**) suggesting that a complex transcriptional program may underlie these subtypes. Integrative analysis comparing the neuron subtypes identified by scRNA-seq and MERFISH (**Fig. 5e** **and Extended Data Fig. 5f-h**) revealed that the majority of D1 and D2 subtypes identified by MERFISH have good correspondence to those identified by scRNA-seq (**Fig. 5f, g** **and Methods**).

**Fig 5.**
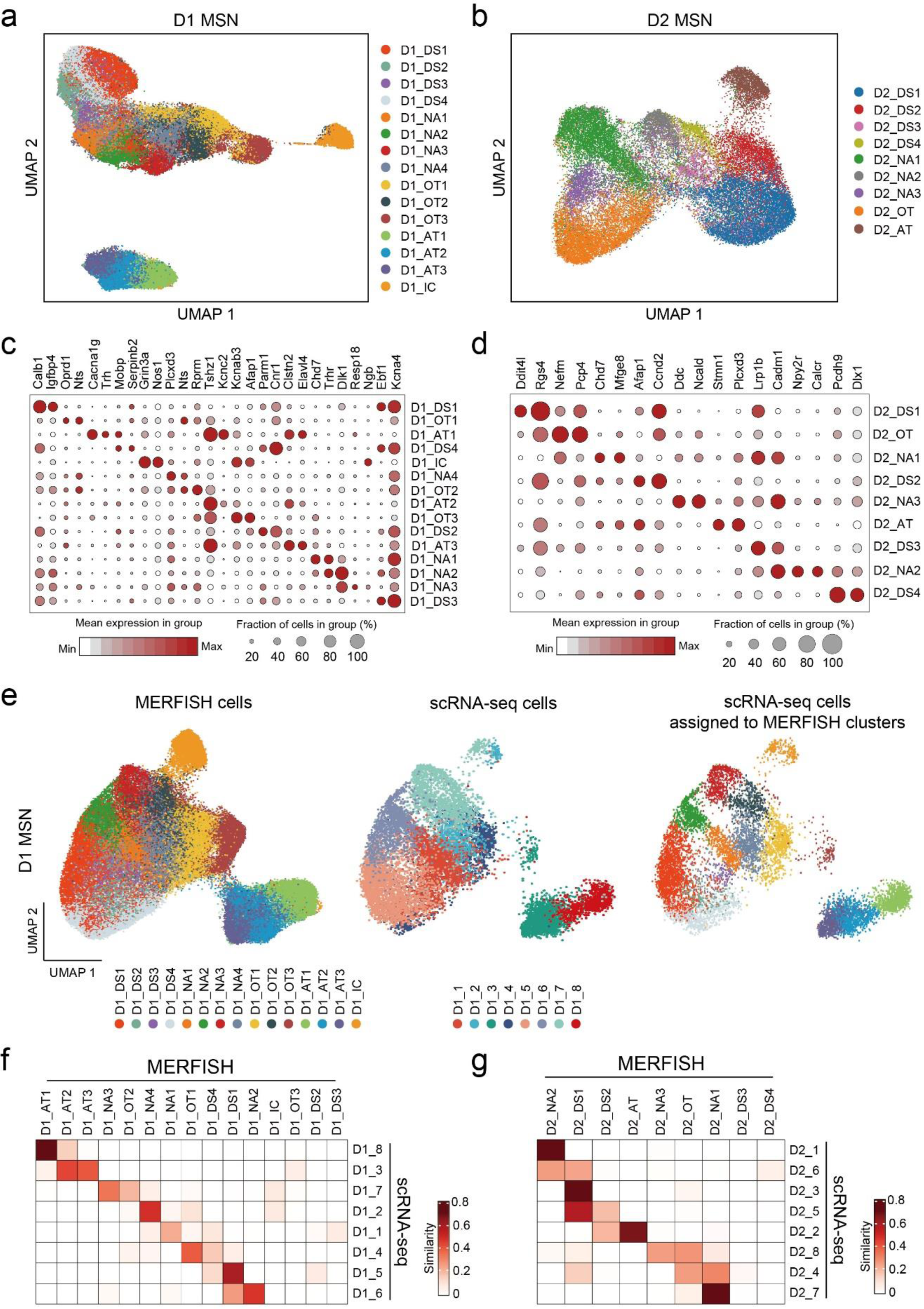
Identification of striatal D1 and D2 MSN subtypes by MERFISH. **a, b,** UMAP plot showing the 15 D1 MSN (a) and 9 D2 MSN subtypes (b) based on MERFISH data. Different MSN subtypes were presented in different colors in the UMAP plot. The subtypes were named based on their spatial distribution. DS, dorsal striatum; NA, nucleus accumbens; OT, olfactory tubercle; AT, atypical; IC, island of Calleja. **c, d,** Dotplot showing the expression pattern of selected genes across the 15 D1 (c) and 9 D2 (d) MSN subtypes. The expression level was color coded. Dot size represents the fraction of cells expressing the gene in each D1 or D2 subtype. **e,** Integrative analysis of D1 MSNs from MERFISH and scRNA-seq experiments. The D1 MSNs from MERFISH (left panel) and scRNA-seq (middle panel) experiments were integrated into the same UMAP space. The identity of each cell was color coded. Based on the nearest neighbors from the MERFISH experiments, the cells from scRNA-seq were assigned to one of the MERFISH D1 MSN subtypes and shown on the right panel. **f, g,** Heatmap showing the correspondence between D1 (**f**) and D2 (**g**) MSN subtypes revealed by MERFISH and scRNA-seq. The MERFISH subtypes without corresponding scRNA-seq subtypes were largely distributed outside of NAc.

Nevertheless, some differences between scRNA-seq and MERFISH subtypes were noted, especially that some neuron subtypes revealed by MERFISH lack a corresponding subtype in the scRNA-seq dataset (**Fig. 5f, g**). A careful analysis of the two datasets revealed that the difference was mainly caused by the different anatomic coverage of the two datasets, as scRNA-seq mainly focused on the NAc while MERFISH analyzed the entire striatum. Because most neuronal subtypes showed a biased distribution in certain striatal subregions (see below), the scRNA-seq data is depleted of the neuron subtypes largely located outside the NAc (**Extended Data Fig. 5h**). For example, the D1_DS2, D1_DS3, D2_DS3 and D2_DS4 subtypes in MERFISH are largely restricted to the dorsal striatum (see below), thus the corresponding subtypes were not identified by scRNA-seq (**Fig. 5f**). Similarly, the MERFISH subtype representing Islands of Calleja (IC) cells (D1_IC) and OT ruffle cells (D1_OT3) were absent in scRNA-seq (**Fig. 5g**) (see below) because they were largely removed during tissue dissection for scRNA-seq. Overall, despite the differences caused by experimental coverage, the neuron subtypes identified by scRNA-seq and MERFISH are largely consistent, which laid a solid foundation for understanding the spatial features of these neuron subtypes.

### Distinct MSN subtypes underlie the structural heterogeneity of the striatum

By plotting MSN subtypes identified by MERFISH as serial striatal sections, we found that different D1 and D2 MSN subtypes exhibit distinct spatial patterns in individual coronal sections (**Fig. 6a, b**). Additionally, the whole dataset spanning >1 mm of brain tissue (from Bregma 1.94 mm to 0.74 mm) enable us to comprehensively assess the MSN subtype distribution along the AP axis. Calculating the proportion of each MSN subtype across all sections revealed that most MSN subtypes exhibit a biased distribution, with different subtypes enriched in different AP coordinates within the striatum (**Fig. 6c, d**). This pattern is in sharp contrast to the largely even distribution of D1 and D2 MSN populations as a whole along the AP axis (**Fig. 4g**), suggesting that the anatomic heterogeneity of striatum is, at least in part, caused by the differential spatial distribution of different MSN subtypes.

**Fig. 6.**
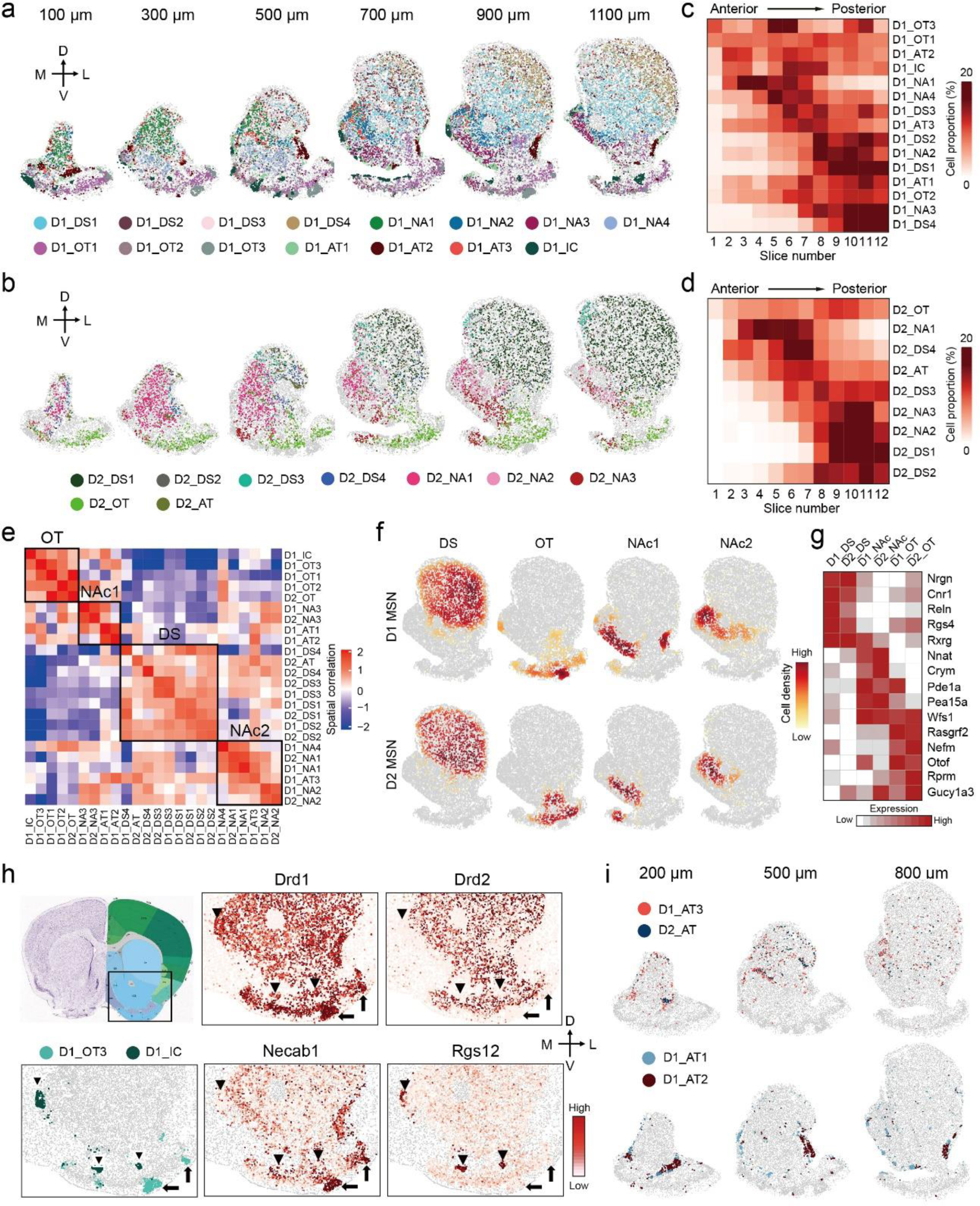
The molecular and spatial features of MSN subtypes underlie anatomic organization of striatum. **a, b,** Spatial patterns of different D1 MSN subtypes (a) and D2 MSN subtypes (b) in coronal sections at different anterior-posterior positions. Six of the twelve slices from a male mouse were shown. Different subtypes were presented by different colors. The 100, 300, 500, 700, 900 and 1100 μm labels indicate the distance from the anterior position (Bregma 1.94mm). The dorsal ventral (DV) and medial-lateral (ML) axes were illustrated. **c, d,** Heatmaps showing the proportion of each D1 MSN (c) and D2 MSN (d) subtype across the 12 coronal sections along the anterior-posterior (AP) axis. **e,** Heatmap showing the spatial correlation across different D1 and D2 MSN subtypes. The black boxes illustrated cell types located in different major striatal divisions, including dorsal striatum (DS), olfactory tubercle (OT) and nucleus accumbens (NAc). **f,** Density maps showing spatial enrichment of different subtype groups of D1 and D2 MSNs in striatal subregions. The MSN subtype groups (DS, OT, NAc1 and NAc2) were the same as (E). The patterns of D1 and D2 MSNs were shown in upper and lower panels, respectively. **g,** Heatmap showing the expression patterns of selected genes in different subtype groups of D1 and D2 MSNs located in major striatal divisions. The subtype groups corresponding to different anatomic regions were determined according to (E), but the two NAc groups were combined. **h,** Specific D1 MSN subtypes represent OT ruffle and IC. The boxed region in the diagram indicates the regions shown in other panels. The OT ruffle and IC were indicated by arrows and arrowheads, respectively. The spatial pattern of D1_OT3 and D1_IC were shown in the lower left panels. The heatmaps showing the expression of *Drd1*, *Drd2*, *Necab1* (marker for D1_OT3) and *Rgs12* (marker for D1_IC) in the corresponding region. **i,** Spatial patterns of three atypical D1 MSN subtypes and one atypical D2 MSN subtype in striatum at different anterior-posterior positions. Different subtypes were presented by different colors. MSN subtypes with higher spatial correlation were shown together. The 200, 500 and 800 μm labels indicate the distance from the anterior position (Bregma 1.94mm).

To understand whether the distribution of MSN subtypes correlates with anatomic organization of striatum, we assessed the spatial colocalization of different MSN subtypes (see Methods). We found that different MSN subtypes are grouped into several “blocks”: the subtypes within the same “blocks” tend to be spatially closer to each other, while subtypes from different “blocks” tend to be spatially separated (**Fig. 6e**). Analysis of the spatial patterns of MSN subtypes in different “blocks” indicated that they represent major anatomic divisions of striatum (**Fig. 6f** **and Extended Data Fig. 6a**). Specifically, D1_DS1 – D1_DS4, D2_DS1 – D2_DS4 and D2_AT form the DS “block” (**Fig. 6e**) representing the dorsal striatum (**Fig. 6f** **and Extended Data Fig. 6a**); the OT “block” containing D1_OT1 – D1_OT3, D1_IC and D2_OT (**Fig. 6e**) corresponds to the OT region (**Fig. 6f** **and Extended Data Fig. 6a**). Interestingly, the rest of the MSN subtypes separated into two groups, with the NAc1 group (D1_NA1, D1_NA2, D1_NA4, D1_AT3, D2_NA1 and D2_NA2) closer to the OT “block” and the NAc2 group (D1_NA3, D1_AT1, D1_AT2 and D2_NA3) closer to the DS “block” (**Fig. 6e**). Both of them represented MSN subtypes in the NAc, but enriched in different subregions (**Fig. 6f** **and Extended Data Fig. 6a**). These results demonstrated that the spatial patterns of certain MSN subtype groups closely resemble major striatal subregions. By further comparing the gene expression of MSN subtypes belonging to different ‘blocks’, we identified genes enriched in the three major subregions: dorsal striatum, NAc and OT (**Extended Data Fig. 6b, c and Supplementary Table 5**). Many of these genes are shared between D1 and D2 MSN subtypes (**Fig. 6g**), suggesting that a global molecular program underlies major striatal anatomic divisions.

In addition to resolving major striatal subregions (dorsal striatum, NAc and OT), the MSN subtypes further illustrate fine anatomic features within these subregions. For instance, the spatial and gene expression features discriminate the D1 MSN subtypes that represent the striosome (D1_DS2) and matrix (D1_DS1, D1_DS3, D1_DS4) compartments ^25^ in the dorsal striatum (**Extended Data Fig. 6d, e**). Additionally, the D1_DS4, D1_DS1 and D1_DS3 form a medial-to-lateral pattern within the dorsal striatum (**Extended Data Fig. 6a**), consistent with the established anatomic and functional distinction along the medial lateral (ML) axis ^43, 44^. Similarly, we found D2 MSN subtypes representing matrix (D2_DS1, D2_DS3)/striosome (D2_DS2) structure (**Extended Data Fig. 6e**) and medial (D2_DS3)/lateral (D2_DS1) striatum (**Extended Data Fig. 6a**). We noted that the majority of D1 MSN subtypes have corresponding D2 MSN subtypes that exhibit very similar spatial patterning (**Fig. 6e** and **Extended Data Fig. 6a**). However, there are some D1 MSN subtypes lacking parallel D2 MSN subtypes, which usually form specific anatomic structures. For instance, the D1_OT1 and D1_OT2 are distributed in the OT flat, with corresponding D2 MSN subtype D2_OT occupying a similar region (**Extended Data Fig. 6a**). In contrast, the D1_OT3 subtype represents the OT ruffle structure and contains only D1 MSNs, without a corresponding D2 MSN subtype (**Fig. 6h**, arrows). Similarly, the Islands of Calleja (IC) is exclusively composed of the D1 MSN subtype D1_IC, which also lacks a parallel D2 subtype (**Fig. 6h**, arrowhead).

Consistent with scRNA-seq results (**Fig. 3d** **and Extended Data Fig. 3b**), MERFISH also revealed a group of atypical MSN subtypes (D1_AT1, D1_AT2, D1_AT3 and D2_AT) that exhibit distinct transcriptional and spatial features (**Fig. 5a, b**). Comparing to the MSNs of the other subtypes, *Tshz1* and *Foxp2* were enriched in the three atypical D1 MSN subtypes, while *Th* and *Tac1* was enriched in D2_AT (**Extended Data Fig. 6f**). Based on their gene expression features, these atypical MSN subtypes correspond to the recently reported eSPN/D1-H SPN (D1_AT1 – D1_AT3) and patch-like D2H SPN (D2_AT) ^29–31^. Our analysis extends previous findings by resolving three subtypes of atypical D1 MSNs, which exhibit distinct transcriptional and spatial features (**Fig. 6i, Extended Data Fig. 6g** and **supplementary Table 6**). Specifically, the D1_AT3 cells (*Tshz1*^+^/*Spon1*^+^) aggregate into a patch-like structure along the border of the anterior striatum, which gradually decreases and shifts to the dorsomedial NAc along the AP axis, with more and more non-aggregating D1_AT3 cells scattered in both dorsal and ventral striatum (**Fig. 6i**). On the other hand, the majority of D1_AT2 cells (*Tshz1*^+^/*Pdyn*^+^/*Penk*^+^) form a dense patch structure at the ventrolateral corner of the NAc, while most D1_AT1 cells (*Tshz1*^+^/*Cacna1g*^+^) form cell clusters along the border of the NAc (**Fig. 6i**). Interestingly, D2_AT and D1_AT3 show similar distribution patterns (**Fig. 6i**), suggesting that they are corresponding MSN subtypes (like other D1/D2 pairs described above). On the other hand, D1_AT1 and D1_AT2 exhibit high spatial correlation (**Fig. 6e, i**) but without any corresponding D2 MSN subtypes. The transcriptional and spatial features of these atypical MSN subtypes suggest that they might be involved in distinct functions that await further investigation ^45^. Collectively, these results suggest that the spatial distribution of distinct MSN subtypes underlie the anatomic complexity of the striatum.

### Substantial MSN heterogeneity in the NAc along the AP axis

Several lines of evidence suggest that compared to the dorsal striatum and OT MSN subtypes, the MSN subtypes in the NAc exhibit more diverse distribution patterns along the AP axes. First, the composition of MSN subtypes comprising the NAc varies from anterior to posterior, as several D1 and D2 MSN subtypes exhibit strong biased distribution along the AP axis. For example, D1_NA1, D1_NA4 and D2_NA1 are enriched in the anterior NAc, while D1_NA2, D1_NA3, D2_NA2 and D2_NA3 are enriched in the posterior NAc (**Fig. 7a, b** **and Extended Data Fig. 7a,b**). Thus, the anterior part of the NAc is dominated by D1_NA1, D1_NA4 and D2_NA1 (**Fig. 7a, b** **and Extended Data Fig. 7a, b**, 100-500 µm slices). Toward the posterior NAc, a new set of MSN subtypes (D1_NA2, D1_NA3, D2_NA2 and D2_NA3) emerge and increase (**Fig. 7a, b** **and Extended Data Fig. 7a, b**, 700-1100 µm slices). The sharp contrast of MSN subtype composition between the anterior and posterior NAc aligns with the anatomic and functional heterogeneity along the AP axis ^6^.

**Fig. 7.**
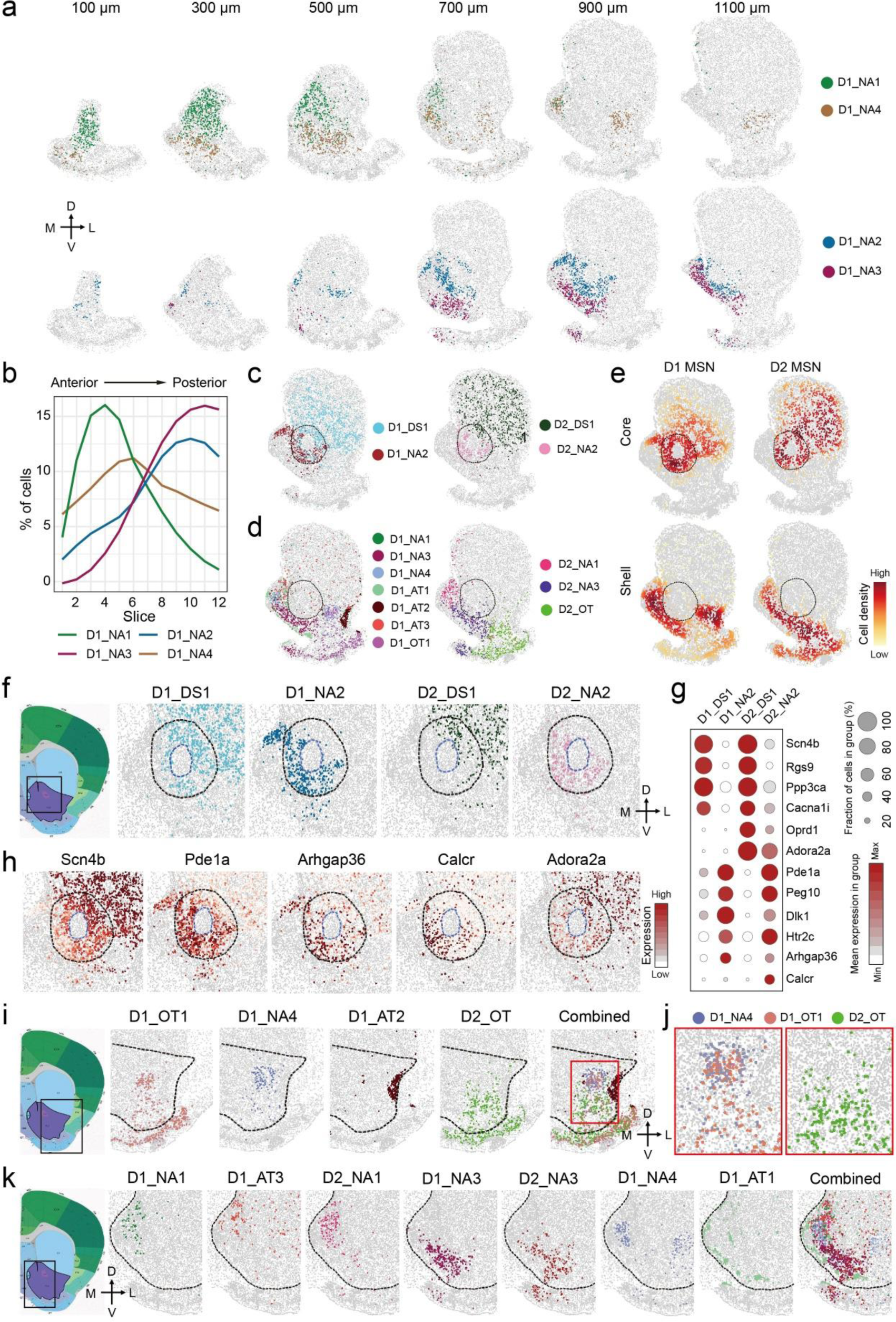
The molecular and spatial features of MSN subtypes underlie anatomic heterogeneity of NAc. **a,** Spatial patterns of selected D1 MSN subtypes in coronal sections of different anterior posterior positions. The subtypes enriched in anterior and posterior NAc were shown in upper and lower panels, respectively. Different subtypes were presented by different colors. The 100, 300, 500, 700, 900 and 1100 μm labels indicate the distance from the anterior position (Bregma 1.94mm). The dorsal-ventral (DV) and medial-lateral (ML) axes were illustrated. **b,** The distribution of the four D1 MSN subtypes shown in (A) along the AP axis. **c,** The spatial pattern of D1 (left panel) and D2 (right panel) MSN subtypes enriched in NAc core. Different subtypes were shown in different colors. The dashed line indicated NAc core. **d,** The spatial pattern of D1 (left panel) and D2 (right panel) MSN subtypes enriched in NAc shell. Different subtypes were shown in different colors. **e,** Density maps showing the spatial pattern of D1 and D2 MSN subtypes enriched in either core or shell subregion of NAc. The subtypes enriched in core and shell subregions of NAc are listed in (c) and (d), respectively. The core region was labeled with dashed line. **f,** Spatial pattern of different MSN subtypes in NAc core. The boxed region in the diagram indicates the regions analyzed in the other panels. The NAc core and AC structure were indicated with dashed lines. The dorsal-ventral (DV) and medial-lateral (ML) axes were indicated. **g,** Dotplot showing the expression of selected gene across D1 and D2 MSN subtypes enriched in different subregions of NAc core. The expression level was color coded. Dot size represents the fraction of cells expressing the gene in each subtype. **h,** Heatmap showing the differential expression of *Scn4b, Pde1a, Arhgap36, Calcr* and *Adora2a* between the dorsolateral and ventromedial part of NAc core. The same region as (F) were shown. The gene expression level was color coded. **i,** Spatial patterns of different MSN subtypes in the lateral shell of NAc. The boxed region in the diagram was shown in the other panels. Different MSN subtypes were color coded. The dashed line represents the border of NAc. The dorsal-ventral (DV) and medial-lateral (ML) axes were indicated. **j,** The red box region in (I) was enlarged to show the spatial distribution of D1_NA4, D1_OT1 and D2_OT in the dorsal part of NAc lateral shell. **k,** Spatial patterns of different MSN subtypes in the medial and ventral shell of NAc. The boxed region in the left panel was analyzed in the other panels. Different MSN subtypes were color coded. The dashed line represents the border of NAc.

Second, the spatial organization of MSN subtypes within the NAc varies from anterior to posterior. In the anterior NAc, the MSN subtypes either form a dorsal-ventral pattern (D1_NA1 and D1_NA4) (**Fig. 7a**) or a largely homogenous distribution (D2_NA1) (**Extended Data Fig. 7a**). On the other hand, in the posterior NAc, the MSN subtypes do not simply follow the dorsal ventral or medial-lateral pattern, but overall form a core/shell-like structure. Currently a core/shell organization of the rodent NAc has been widely accepted, but the underlying cellular basis remains unclear. Analysis of our MERFISH data revealed that in the posterior NAc (700-1100 µm slices), four major MSN subtypes (D1_DS1, D1_NA2, D2_DS1 and D2_NA2) occupy the region surrounding the AC, which is considered as the NAc core (**Fig. 7c** **and Extended Data Fig. 7c**). Other MSN subtypes in the NAc are located largely surrounding this region, which is considered as the NAc shell (**Fig. 7d** **and Extended Data Fig. 7c**). The density map of these different MSN groups clearly illustrated the core/shell subregions (**Fig. 7e**). Notably, this MERFISH defined core/shell pattern could not be simply recapitulated by the expression pattern of any single gene, although some genes (eg. *Calb1* and *Gucy1a3*) are highly enriched in certain MSN subtypes located in the core/shell region (**Extended Data Fig 7d**). This indicates that complex transcriptional programs of multiple neuron subtypes underlie the core/shell organization. On the other hand, although certain MSN subtypes are largely restricted to either the core or the shell of the NAc, they do occupy overlapping regions (**Extended Data Fig. 7c**). This may explain why previous studies using limited marker genes could not identify a definitive border of the core/shell subregions ^12, 24^. Together, our analyses reveal substantial neural heterogeneity along the AP axis of the NAc and suggest that spatial enrichment of specific MSN subtypes could be an important factor underlying the anatomic features of the NAc, including the core/shell division.

### Substantial MSN subtype heterogeneity within the core/shell subregions of the NAc

In addition to the general differences between the NAc core and shell described above, analyses of the MERFISH dataset have uncovered previously unrecognized heterogeneity within the core and shell subregions. In the NAc core, MERFISH revealed a ventromedial-dorsolateral patten spanning this region. Specifically, the D1_NA2 and D2_NA2 subtypes are preferentially located in the ventromedial core, while D1_DS1 and D2_DS1 subtypes are enriched in the dorsolateral part of the core (**Fig. 7f** **and Extended Data Fig. 7e**). This pattern suggests that the dorsolateral and ventromedial part of the NAc core possess different features, with the former resembling the anatomy features of the adjacent dorsal striatum ^12^, as D1_DS1 and D2_DS1 are two of the major MSN subtypes in the dorsal striatum (**Fig. 7c** **and Extended Data Fig. 7c**). To gain insights into the ventromedial-dorsolateral pattern of the NAc core at the molecular level, we identified differentially expressed genes (DEGs) between MSN subtypes enriched in the two subregions (**Fig. 7g**, **Extended Data Fig. S7f** **and Supplementary Table 7**). As expected, these DEGs exhibited substantial bias in expression in corresponding subregions of the NAc core (**Fig. 7h** **and Extended Data Fig. 7g**), and many DEGs are shared between D1 and D2 subtypes (**Fig. 7g**), suggesting that a common transcriptional program underlies the cellular/anatomic heterogeneity of the NAc core. Interestingly, we also found some DEGs are specific to D1 or D2 MSNs, such as *Arhgap36* (D1 specific), *Calcr* and *Adora2a* (D2 specific) (**Fig. 7g, h**), suggesting that a cell type-specific mechanism may also be involved. Since quite a few DEGs are pertinent to neuronal function, including genes encoding receptors (*Adora2a*, *Htr2c*, *Calcr*, *Oprd1*), channels (*Cacna1i*, *Scn4b*) and enzymes (*Ppp3ca*, *Pde1a*) (**Fig. 7g, h**), MSN subtypes enriched in different subregions of the NAc core can differentially respond to upstream signals, linking the anatomy of the NAc core to diverse neuronal functions.

Previous studies have revealed prominent anatomic and functional heterogeneity within the NAc shell ^6, 16, 24, 27^. Consistently, we found MSN subtypes within this region form a more complex topographic map compared to the core region. First, there are more D1 and D2 MSN subtypes located in the NAc shell. At least seven D1 (D1_NA1, D1_NA3, D1_NA4, D1_AT1, D1_AT2, D1_AT3, D1_OT1) and three D2 (D2_NA2, D2_NA3 and D2_OT) MSN subtypes have significant distribution within the NAc shell (**Fig. 7d** **and Extended Data Fig. 7c**), compared to only two D1 and two D2 MSN subtypes that are enriched in the NAc core (**Fig. 7c** **and Extended Data Fig. 7c**). In addition, different MSN subtypes exhibit distinct spatial patterns, leading to different neuronal composition in the lateral and medial part of the NAc shell. In the lateral shell, the D1_NA4 and D1_OT1 subtypes are located in its medial part (adjacent to the NAc core), while D1_AT2 forms a dense patch structure in the most lateral region (adjacent to cortex) (**Fig. 7i**). On the other hand, only one dominant D2 MSN subtype, D2_OT, is located in the lateral shell (**Fig. 7i**). Interestingly, the D2_OT subtype is enriched in the ventral part of the lateral shell, but depleted from the dorsal part, resulting in a region largely occupied by D1 MSNs in the dorsolateral shell (**Fig. 7j**). This observation was further supported by MERFISH and ISH detection of D2-receptor mRNA (*Drd2*) in this region (**Extended Data Fig. 7h**). We speculate that the dopamine signaling in this D2 MSN-depleted region is highly biased toward a D1-receptor pathway and is predominantly mediated by D1_NA4 and D1_OT1.

Compared to the lateral shell, the medial shell exhibits even higher heterogeneity in terms of MSN composition and their spatial pattern. At least five D1 (D1_NA1, D1_NA3, D1_NA4, D1_AT1, D1_AT3) and two D2 (D2_NA1 and D2_NA3) MSN subtypes are enriched in the medial shell (**Fig. 7k**). Among these subtypes, D1_NA1, D1_AT3 and D2_NA1 show a similar distribution pattern, with all of them occupying the dorsal part of medial shell (**Fig. 7k** **and Extended Data Fig. 7i**). In contrast, D1_NA3 and D2_NA3 subtypes mainly occupy the ventromedial and ventral part of the shell (**Fig. 7k** **and Extended Data Fig. 7i**). For the other two D1 subtypes, D1_NA4 mainly occupies the middle between D1_NA2 and D1_NA3 (**Fig. 7k** **and Extended Data Fig. 7i**), while D1_AT1 forms small cell clusters along the border of the medial shell (**Fig. 7k**). Despite the distinct spatial patterns, multiple MSN subtypes always co occupy any given subregions in the medial shell (**Fig. 7k**), highlighting the neuronal complexity in this region.

Interestingly, we note that the topological organization of the core and shell subregions described here are mainly observed in coronal sections from the posterior NAc (700-1100 µm slices in our MERFISH dataset), while MSN subtypes and their distribution patterns are substantially different in the anterior NAc (100-500 µm slices in our MERFISH dataset, see above), adding another layer of neuronal complexity to the NAc. Overall, our results reveal a high level of MSN heterogeneity within the core and shell subregions of the NAc. The molecular and spatial features of different neuron subtypes revealed by MERFISH are consistent with early anatomic studies based on immunostaining and neural tracing ^12^, supporting the notion that neuronal diversity underlies the anatomic heterogeneity of the NAc.

### Spatial features of MSN subtypes could link MSN subtypes to specific neuronal functions

Having established that the spatial pattern of molecularly defined MSN subtypes underlies the anatomic heterogeneity of the NAc, we next asked whether the molecular and spatial features of these neural subtypes are related to the functional complexity of the NAc. Previous studies have revealed a functional difference between the NAc core and shell ^46–48^. By revealing the distinct MSN subtypes in these subregions (**Fig. 7c-e**), we can now link previously identified core- or shell-specific functions to specific MSN subgroups located in the corresponding region. Furthermore, our data revealed further spatial heterogeneity within NAc subregions, which is also well aligned with functional heterogeneity observed in the NAc. For example, previous studies have showed that applying opioid receptor ligands to the medial NAc along the AP axis have different impact on hedonic response (“hot spot” and “cold spot”) ^49^. This functional change is consistent with the observed changes of MSN subtypes along the AP axis (**Fig. 7a, b** **and Extended Data Fig. 7a, b**). Similarly, the functional difference of Pdyn^+^ (a pan D1 MSN marker in the NAc) MSNs that reside in the dorsomedial and ventromedial NAc ^27^ could also be explained by MSN subtype-specific function, as distinct *Pdyn*^+^ MSN (D1 MSN) subtypes are enriched in these subregions (**Fig. 7k** **and Extended Data Fig. 7i**). Interestingly, a recent study revealed that dopamine signaling in the ventromedial NAc specifically encodes aversive stimuli ^17^, and this same region is occupied by D1_NA3 and D2_NA3 (**Fig. 7k** **and Extended Data Fig. 7i**), suggesting that these two MSN subtypes can potentially serve as the neuronal substrates of this region-specific dopamine signal. Furthermore, another recent study revealed an unexpected role of the Tshz1^+^/Pdyn^-^ D1 MSNs (corresponding to the D1_AT3 subtype based on gene expression and spatial distribution) in aversive coding and learning ^45^, providing strong evidence that molecularly defined MSN subtypes are functionally distinct.

### Differential gene expression in different MSN subtypes underlying functional complexity of the NAc

To gain cellular and molecular insights into the functional complexity of the striatum, we assessed the expression of genes relevant to neural function (neural peptide, receptors, ion channels) and found that these genes exhibit very different expression patterns among different MSN subtypes (**Fig. 8a**). For example, *Trhr* (encoding Trh receptor) is mainly expressed in the D1_NA1, D1_NA2, D2_NA1 and D2_NA2 subtypes in the NAc (**Fig. 8a**), suggesting that the role of Trh in the NAc in feeding regulation ^50^ is mediated by these MSN subtypes. On the other hand, *Oxtr* (encoding oxytocin receptor), *Drd3* (encoding dopamine receptor 3) and *Npy1r* (encoding NPY receptor 1) are highly enriched in cells of the IC (D1_IC) (**Fig. 8a**), suggesting the IC may serve as a hub to integrate divergent upstream signals to regulate behavior.

**Fig. 8.**
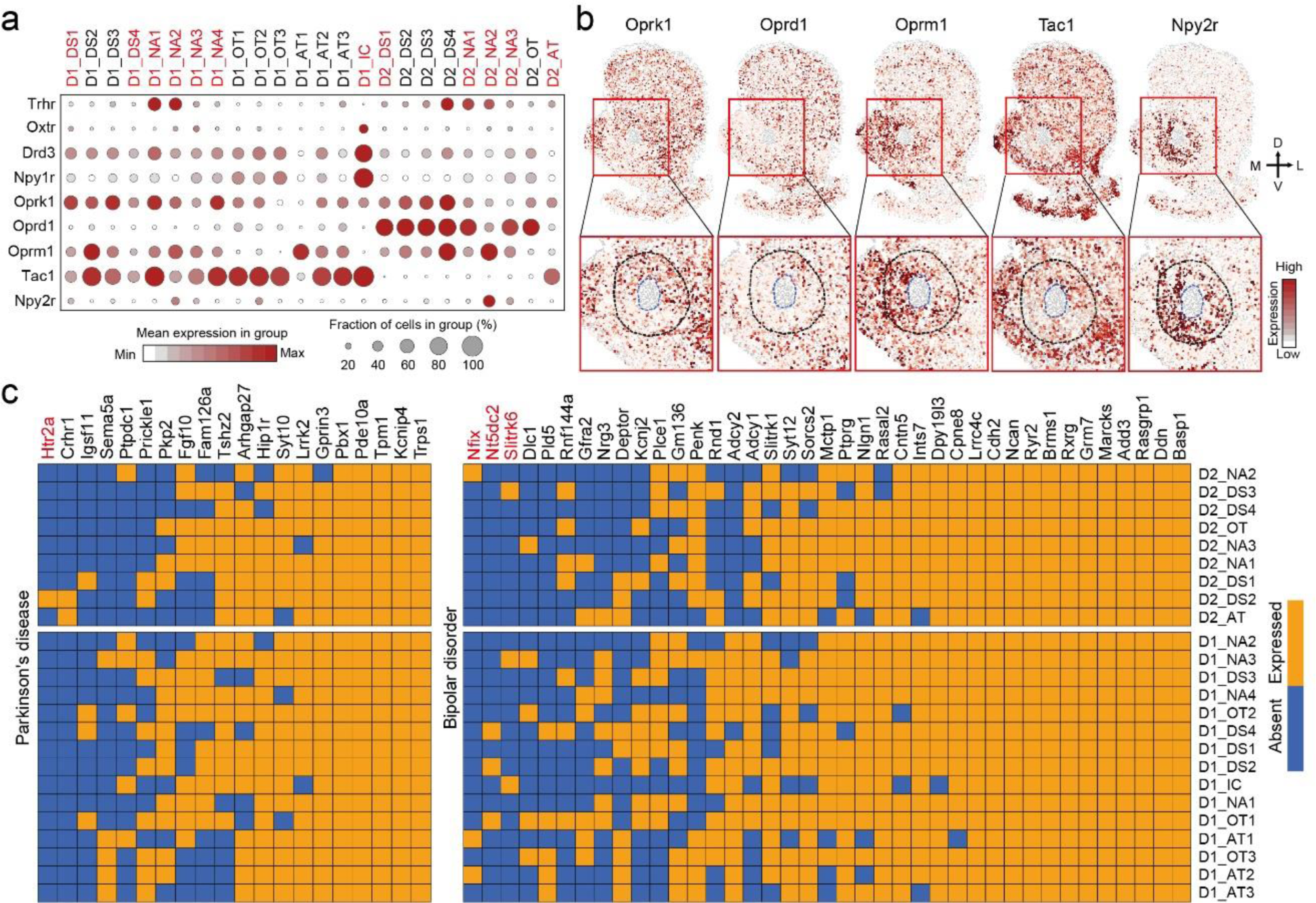
The molecular and spatial features of neuron subtypes underlie functional complexity of NAc. **a,** Dotplot showing the expression patterns of selected functional-relevant genes across all the D1 and D2 MSN subtypes. The expression level is color coded. The MSN subtypes mentioned in the text are labeled in red. **b,** Heatmap showing the expression patterns of *Oprk1*, *Oprd1*, *Oprm1*, *Tac1* and *Npy2r* in striatum. The boxed regions in the upper panels were enlarged and shown in lower panels. The NAc core and AC structure were labeled with dashed lines. Note that all these genes exhibit differential expression between NAc core and shell or between subregions within the core. The gene expression level was color coded. **c,** Heatmap showing the expression pattern of GWAS candidate genes associated with Parkinson’s disease and bipolar disorder across different D1 and D2 MSN subtypes. Blue color indicates no enrichment, orange color indicates gene enrichment. The genes mentioned in the text are labeled in red.

We paid special attention to genes encoding opioid receptors because extensive studies have revealed their complex functions in the striatum ^51, 52^. We found that different opioid receptors exhibit distinct expression patterns (**Fig. 8a, b**). For example, *Oprk1* (encoding κ-opioid receptor) is widely expressed in the majority of D1 and D2 MSN subtypes with significant variation along the AP axis. Specifically, *Oprk1* is enriched in the D1_NA1 and D1_NA2 subtypes, but is depleted in the D1_NA3, D2_NA2 and D2_NA3 subtypes (**Fig. 8a**). Since these two groups of MSNs are respectively biased toward the anterior and posterior part of the NAc (**Fig. 6c, d**), *Oprk1* exhibits a high-to-low expression pattern along the AP axis in the ventromedial NAc (**Extended Data Fig. 8a**). In contrast, *Oprd1* (encoding δ-opioid receptor) is restricted to D2 MSNs and expressed in most D2 subtypes (**Fig. 8a**). However, two of the D2 subtypes are significantly depleted in *Oprd1* expression, including the D2_AT subtype dispersed across the striatum (**Fig. 6i**) and the D2_NA2 subtype located in the ventromedial part of the NAc core (**Fig. 7f**). Because *Oprd1* is highly expressed in the D2_DS1 subtype that occupies the dorsolateral core (**Fig. 7f**), we see substantial differential expression of *Oprd1* in the NAc core along the ventromedial-dorsolateral axis (**Fig. 8b**). Interestingly, we found that *Oprm1* (encoding µ-opioid receptor) forms a complementary spatial pattern to that of *Oprd1*, which is enriched in D1_NA2 and D2_NA2 subtypes, but shows relatively low expression in the D1_DS1, D2_DS1, D1_NA1 and D2_NA1 subtypes (**Fig. 8a, b**). These cell type and region-specific enrichments of different opioid receptors suggest that opioid signaling is involved in complex regulation and interaction of different MSN subtypes ^53, 54^, and likely underlie their functional complexity in different NAc subregions ^6, 49^.

### Spatial and transcriptional information reveal potential subregion and cell type-specific interaction

In addition to inferring the functional properties of individual neuronal subtypes, the spatial and transcriptional information obtained from MERFISH could also be used for identifying potential cell-cell interactions. To illustrate this application, we focused on the neuropeptide-receptor communication between MSN and interneuron subtypes, which suggested potential region and cell type-specific regulatory relationships in the striatum. For example, we found that *Tac1* is expressed in the majority of D1 MSN subtypes. Given that Tac1 receptor gene, *Tacr1*, is expressed in both *Chat*^+^ (cholinergic) and *Npy*^+^ interneurons, a wide-spread interaction between D1 MSN and these two interneuron subtypes through a *Tac1*-*Tacr1* pathway is expected (**Extended Data Fig. 8b**). Interestingly, a few D1 MSN subtypes (D1_NA2, D1_DS1, D1_DS4) showed a significantly lower *Tac1* level (**Fig. 8a**), suggesting that this regulatory pathway is attenuated in the corresponding regions that harbor these subtypes, including the NAc core (**Fig. 8b**) and the matrix in the dorsal striatum. On the other hand, *Npy*^+^ interneuron (**Extended Data Fig. 8b**) may exert a region-specific function in regulating the NAc core through the *Npy*-*Npy2r* pathway, as *Npy2r* is enriched in the MSN subtypes (D1_NA2 and D2_NA2) located in this region (**Fig. 8b**). Notably, by integrating spatial information, the cell-cell communication implied from gene expression could be evaluated more accurately. For instance, we found *Trh* is expressed in *Th*^+^/*Trh*^+^ interneurons (**Extended Data Fig. 8c**) and *Trhr* is expressed in certain MSN subtypes (**Fig. 8a** **and Extended Data Fig. 8c**). However, few *Trh*^+^ interneurons are distributed in the *Trhr*^+^ MSN-enriched region (dorsomedial NAc) (**Extended Data Fig. 8c, arrow head**), suggesting that an additional Trh source, such as *Trh*^+^ neurons in the hypothalamus ^55^, may act on these *Trhr*^+^ MSNs. Although our analysis has focused on genes directly detected by MERFISH, the results support the notion that highly divergent cell-cell interactions in the striatum depend on both transcriptional and spatial features of different neuronal subtypes.

### Neurological disorder-related genes are differentially expressed in MSN subtypes

To gain further insights into how different neuronal subtypes may link to different striatal functions, we extended our analysis to the entire transcriptome by integrating the scRNA-seq and MERFISH dataset (**Fig. 5e** **and Extended Data Fig. 5f**) to infer the expression of genes not targeted in MERFISH from the scRNA-seq data. Using this integrative analysis, we asked whether the different MSN subtypes may be associated with specific brain disorders. To this end, we assessed whether candidate genes identified from genome wide association studies (GWAS) of different neural disorders were enriched or selectively expressed in certain MSN subtypes. We chose eight neural disorders that have been linked to striatal dysfunction, including schizophrenia, bipolar disorder, depression, Parkinson’s disease (PD), obsessive compulsive disorder, alcohol addiction and nicotine addiction, and obtained GWAS candidate genes previously linked to these disorders from the EMBL GWAS catalog. By assessing the expression profile of these GWAS candidate genes in different MSN subtypes, we found that the majority of these genes are widely expressed across all MSN subtypes (**Fig. 8c** **and Extended Data Fig. 8d**), suggesting that dysfunction of these genes may have a broad impact on striatal function. However, a significant proportion of these genes (25-50%) exhibit selective/enriched expression in less than 50% of all MSN subtypes (**Fig. 8c** **and Extended Data Fig. 8d**). For example, *Htr2a*, a PD associated gene encoding the serotonin receptor, is selectively expressed in the D2_DS2 subtype (**Fig. 8c**). On the other hand, three bipolar disorder-associated genes, *Nfix*, *Nt5dc2* and *Slitrk6*, are expressed in non-overlapping D1 and D2 MSN subtypes (**Fig. 8c**), suggesting that a cell type-specific mechanism potentially contribute to the pathogenesis of this disease. Similar examples can be found in all other neural disorders analyzed (**Extended Data Fig. 8d**). Collectively, these results suggest the possibility that cell type-specific transcriptional programs could render different vulnerability to striatal neuron subtypes in the pathogenesis of brain disorders.

## Discussion

Single-cell transcriptomic profiling has greatly advanced our understanding about the cellular heterogeneity of the nervous system ^56^. However, most high-throughput scRNA-seq methods use dissociated cells and thus do not provide spatial information of analyzed cells, which makes it challenging to connect the molecular feature of the relevant neuron types to their anatomic and functional features. Although this problem can be partially addressed by linking the neuron subtype marker gene expression to known spatial features (such as cortical layers) in brain regions with clear spatial organization ^57–59^, the same strategy is difficult to apply to regions lacking clear anatomic features, such as the NAc. By carefully analyzing the single-cell transcriptional program in the striatum, a recent study revealed that both discrete and continuous gene expression features underlie the MSN heterogeneity, which is closely related to their spatial organization in striatum ^30^. A systematic and high-resolution characterization of the spatial organizations of striatal neuronal subtypes remains highly desirable to bridge the molecular and spatial features of cell types with the functional heterogeneity of the NAc. The rapid development of imaging-based tools in the past several years has enabled in situ, spatially resolved, single-cell transcriptomics analyses ^60^. By combining two complementary methods, scRNA-seq and MERFISH, here we generated a NAc taxonomy with both transcriptional and spatial information at single-cell resolution. Our study not only revealed diverse neuron subtypes with distinct molecular and spatial features, but also linked the gene expression and spatial distribution of these neuron subtypes to the anatomic and functional complexity of the NAc, thus providing insights into how the NAc, which has a seemingly simple neuronal composition (D1 and D2 MSNs), accomplishes its observed structural and functional diversity.

### Transcriptional and spatial differences of MSN subtypes underlie NAc anatomic organization

The anatomic heterogeneity of the NAc, especially the core/shell division, has been recognized for decades ^24^. The core/shell subregions were initially proposed based on differential expression of several marker genes and the notion was further supported by studies illustrating anatomic and functional distinction ^12, 47^. However, the basis of these anatomic structures remains unclear, and definitive criteria distinguishing the NAc core and shell is lacking. By assessing the spatial organization of transcriptionally distinct MSN subtypes, we have identified a cellular basis for the core/shell subregions, and uncovered novel anatomic complexity beyond these known NAc subregions. First, the anterior and posterior part of the NAc exhibited substantial difference in the composition of MSN subtypes as well as spatial organization, with the core/shell structure only prominent in the posterior part. Second, the core/shell division is resembled by the spatial pattern of two groups of MSN subtypes. Third, further anatomic complexity, characterized by the spatial organization of different MSN subtypes, was noticeable even within the core and shell region. Fourth, despite that most MSN subtypes show strong bias to certain NAc subregions, there is often spatial overlap among different MSN subtypes, which explains the absence of a sharp anatomic border separating these subregions. Notably, a recent study suggested that continuous transcriptional pattern underlying the spatial organization of striatum ^30^. Consistent with this notion, we detected similar gradient expression of certain genes across MSN subtypes and large striatal regions (e.g. **Extended Data Fig. 6c**). Furthermore, with the power of detecting hundreds of genes at single-cell resolution *in situ*, we identified more diverse and complex transcriptional patterns at higher resolution, especially in NAc (e.g. **Fig. 7h**), suggesting a tight correlation between gene expression and spatial features of MSNs at different scales. It is also worth mentioning that several recent studies have revealed distinct neural connections that originate from or project to different NAc subregions ^16, 61–63^, which are consistent with the spatial distribution of certain neuron subtypes described in this study. All these findings support the notion that molecularly defined neuron subtype and their spatial distribution is a fundamental factor underlying the anatomic organization of the NAc.

### Neuronal heterogeneity underlies functional complexity of the NAc

A large body of work has revealed the functional diversity of the NAc ^1^ and its dysfunctions have been linked to multiple neural disorders ^7–10^. This functional diversity cannot be well explained by the conventional direct/indirect model and is usually associated with different subregions of the NAc. Indeed, intra-NAc functional heterogeneity along multiple anatomic axes has been described ^17, 49^, and some recent studies have demonstrated region-specific functions by directly manipulating neurons located in different NAc subregions ^16, 27^. However, the general mechanism underlying this regional-specific function is unclear. Our work helped fill in this knowledge gap by linking functional diversity of the NAc to its different neuronal subtypes and their distinct molecular and spatial features. Specifically, we revealed: 1) the spatial distribution of MSN subtypes at single-cell resolution, providing a framework for mapping previously reported functional variation along the AP, DV and ML axes in the NAc; 2) variable expression of neural function-relevant genes among MSN subtypes, leading to cell-type- and region-specific expression patterns that likely confer region-specific function; 3) potential cell-type- and region specific interactions within the NAc, leading to a complex regulatory network; 4) genome-wide transcriptional profiles of different neuronal subtypes (by integration of scRNA-seq and MERFISH data), which could potentially be associated with different neural disorders. Interestingly, a recent study assessing the Tshz1^+^ neurons in the dorsal striatum ^45^ provided direct evidence that molecularly defined MSN subtype can possess distinct function. It is worth mentioning that our data is not contradictory to the conventional dichotomy model, as most NAc subregions were well covered by both D1 and D2 MSNs, and the majority of D1 MSN subtypes have corresponding D2 subtypes that show similar spatial distribution patterns. These data suggest that the dichotomy model is recapitulated on the sub-region scale and that D1/D2 MSNs work in concert as fundamental components of NAc computation. Together, by identifying the molecular and spatial patterns of different neuron subtypes, our study has established a connection between neuron diversity and functional heterogeneity of the NAc.

### A framework for understanding the anatomic and function of the NAc

In the cerebral cortex, the relationship between molecularly defined cell types and anatomic organization has been extensively studied ^57, 58^, which facilitated cell type-specific functional characterization ^64–66^. Here we provide evidence suggesting a similar molecular-cellular anatomic-functional relationship exists in the NAc, but in a more complex fashion. Thus, our transcriptomic-based NAc cell taxonomy can serve as a framework to help further our understanding of the structural and functional relationship of the NAc in multiple ways. First, identification of transcriptionally different neuron subtypes in the NAc will facilitate the development of new genetic tools, including transgenic animals and viruses. Since most NAc subregions are occupied by multiple neuronal subtypes, genetic approaches are essential for dissecting cell type-specific roles in health and disease. Second, the cell taxonomy could be used to annotate and infer neuronal substrates if genetic/molecular information (eg. transgenic line or marker gene staining) is available, making direct comparison of different studies possible. This is especially important when similar or adjacent NAc subregions are targeted. Third, the MERFISH and scRNA-seq approaches used in this study can be combined with neural tracing and activity mapping ^33, 67^ to further integrate anatomic and functional information in understanding neuronal processes. Notably, recent studies have comprehensively mapped the region-specific inputs and outputs of the dorsal striatum ^43, 68, 69^. Combining a similar approach with MERFISH could potentially reveal the relationship between molecularly defined MSN subtypes and the region specific afferent/efferent of the NAc. Fourth, in addition to linking known anatomic and functional heterogeneity to different neuronal subtypes, our transcriptional and spatial analyses also revealed substantial uncharacterized molecular and cellular heterogeneity in the NAc (such as the ventromedial-dorsolateral difference in the NAc core), which can serve as the basis for generating novel hypotheses for further studies.

### Data visualization and sharing

To maximize the usage of the large datasets generated in this study, and to help people easily explore the datasets, we have setup an interactive website-based tool to visualize our scRNA-seq (http://35.184.4.122:9090/) and MERFISH (http://35.184.4.122:5050/) data.

## Materials and Methods

### Mice

All experiments were conducted in accordance with the National Institute of Health Guide for Care and Use of Laboratory Animals and approved by the Institutional Animal Care and Use Committee (IACUC) of Boston Children’s Hospital and Harvard Medical School. For single-cell RNA-seq, we used 10 weeks young adult male C57BL/6N mice (Cat# 000664, Jackson Lab) and the NAc tissue were collected for single-cell RNA-seq. The mice were housed in groups (3-5 mice/cage) in a 12-hr light/dark cycle, with food and water *ad libitum*. For FISH and MERFISH assays, we used 8-10 week-old male C57BL/6N mice.

### Tissue dissection and dissociation

Dissection and cell dissociation were performed in 11 separate experiments, with brain tissues from two mice were pooled for cell dissociation and library preparation in each experiment. Each experiment was regarded as one biological replicate. For single cell dissociation of nucleus accumbens (NAc), the mice were anesthetized with isoflurane and the brain was quickly removed and transferred into ice-cold Hibernate A/B27 medium (60 ml Hibernate A medium with 1 ml B27 and 0.15 ml Glutamax). Coronal sections containing NAc were cut using a brain matrix and sliced into 0.5 mm slices in ice-cold Hibernate A/B27 medium. NAc tissue was then removed from each slice under dissection microscope and subjected to tissue dissociation. NAc tissues were further dissociated into single-cell suspension using a papain-based dissociation protocol ^70^ with some modifications. Briefly, the tissues from two animals were cut into small pieces and incubated in dissociation medium (Hibernate A-Ca medium with 2 mg/ml papain and 2X Glutamax) at 30 °C for 35-40 min with constant agitation. After washing with 5 ml Hibernate A/B27 medium, the tissues were triturated with fire polished glass Pasteur pipettes 10 times in 2 ml Hibernate A/B27 medium to generate single-cell suspension, which was repeated 3 times. To remove debris, the 6 ml single-cell suspension was loaded on a 4-layer OptiPrep gradient ^70^ and centrifuged at 800 g for 15 min at 4 °C. Fractions 2 – 4 were then collected and washed with 5 ml Hibernate A/B27 medium and 5 ml DPBS with 0.01% BSA. The cells were spun down at 200 g for 3 min and re-suspended in 0.2 ml DPBS with 0.01% BSA. A 10 μl cell suspension was stained with Trypan Blue and the live cells were counted. During the entire procedure, the tissues and cells were kept in ice-cold solutions except for the papain digestion.

### Single cell capture, library preparation, and sequencing

The cells suspension was diluted with DPBS containing 0.01% BSA to 300 - 330 cells/μl for single cell capture. Single cells and barcoded beads were captured into droplets with the 10X Chromium platform (10X Genomics, CA) according to the protocol from the manufacturer ^71^. After cell capture, reverse transcription, cDNA amplification and sequencing library preparation were perform as described previously ^71^. The libraries were sequenced on Illumina Hiseq 2500 sequencer with pair-end sequencing (Read1: 26bp, Index: 8bp, Read2: 98bp).

### Fluorescence in situ hybridization (RNAscope) and imaging

For sample preparation, young adult male (8 - 10 weeks) C57B6 mice (Jackson Lab) were anesthetized and perfused with PBS followed by 4% paraformaldehyde (PFA) in PBS. The whole brain was dissected and post-fixed in 4% PFA overnight, followed by dehydration in PBS containing 30% sucrose at 4 °C until the tissues sank to the bottom of the tube. The brains were frozen in Optimal Cutting Temperature (OCT) embedding media and 20 μm (for FISH) or 40 μm (for immunostaining) coronal sections were cut with cryostat. For FISH, the slices were mounted on SuperFrost Plus slides, air dried and store at -80 °C until use. For immunostaining, the slices were stored in PBS at 4 °C until use. The multi-color FISH experiments were performed following the manual of ACD RNAscope Fluorescent Multiplex Assay. For imaging, brain sections were imaged on a Zeiss confocal microscope (LSM800) with a 10x (0.3 NA) or 20x (0.8NA) objective. Z stacks were taken with 1 μm optical sectioning. For some sections, tiled images were acquired which covered the whole NAc region.

### Generation of single-cell gene expression matrix

Raw sequencing data were processed with cellranger (v 1.3.1) ^71^ for sample demultiplexing, cell barcode detection and single-cell expression matrices generation. Briefly, the “cellranger mkfastq” command was used to demultiplex the different samples, extract the UMI barcodes and generate the fastq files. The gene-cell expression matrices for each sample were generated separately using the “cellranger count” command by aligning the reads to the mm9 genome. In total, 11 single-cell gene expression matrices corresponding to the 11 biological replicates were generated (Saline: 2 samples in Maintenance phase, 1 sample 48h withdrawal phase, 2 samples 15 days withdrawal phase. Cocaine: 2 samples Maintenance phase, 2 samples withdrawal phase, 2 samples 15 days withdrawal phase).

### Single-cell transcriptomic data filtering and quality control

The R package Seurat (v2.1.0) ^72^ and some customized R scripts were used for data analyses. First, the expression matrices of all the 11 IVSA samples under saline and cocaine treatments were pooled together into a global Seurat object using the “MergeSeurat” function. Genes expressed in less than 3 cells were excluded. The cells with >10% of their total transcripts from mitochondrial transcriptome and cells with a large number of uniquely expressed genes (more than the 99th percentile, 4766 genes) were removed. For the initial analysis, all single cells expressing ≤ 1500 genes were also excluded. In total, an expression matrix including 19,458 genes and 37,011 cells were obtained.

The gene expression profile of each single cell was then normalized to count per-million (cpm) and natural log transformed. To further exclude potential experimental variations, a linear model was generated by using the ScaleData function in Seurat to regress out the effect of the 1) number of detected UMI in each cell, 2) percent of mitochondrial genes and 3) sample-to-sample variation.

### Separation of cells into neuronal and non-neuronal populations and detection of major cell populations

To detect the major neuronal and non-neuronal populations, 2,104 genes with cpm > 1.5 and dispersion (variance/mean expression) larger than one standard deviation away from the expected dispersion were selected as variable genes using the Seurat function “FindVariableGenes”. The variable genes were used to calculate the top 30 principle components (PCs) and the significant PCs were selected using the “Jackstraw” method in the Seurat package (p-value < 1e-3). The cells were then classified into 25 clusters with the Seurat “FindClusters” function. Then, the 25 initial clusters were merged using a random forest classifier with 10-fold cross validation and 500 trees built with the caret R package ^73^, until the markers of each cluster having a strong predictive power (>80%), which led to 13 cell clusters. Using the saline treated samples and based on the expression of neuron-specific gene Snap25, the 13 cell clusters were identified as neuronal or non-neuronal clusters. In total, 5 neuronal clusters with 26,429 cells and 8 non-neuronal clusters with 10,582 cells were detected. These 13 major cell clusters were further aggregated into 5 non-neuronal populations (astrocyte, endothelial cells, microglia, oligodendrocyte and oligodendrocyte precursor cells) and 4 neuronal populations (D1 MSN, D2 MSN, inter-neuron and new-born neuron) based on the expression of established marker genes, and is presented in Figure 1.

### Inclusion of non-neuronal cells with > 800 genes detected

Previous studies revealed that neuronal cells tend to have a higher number of mRNAs (UMI) and genes than non-neuronal cells ^34, 35^, which is also confirmed in our dataset (**Fig. S1D**). Thus, the criteria used above (>1500 gene detected) excluded many non-neuronal cells. Hence, we decided to include non-neuronal cells with >800 detected genes, which was regarded as high-quality cells in previous high-throughput single-cell RNA-seq studies ^74, 75^. To this end, the cells expressing ≥ 1500 genes were used to train a random forest classifier (500 trees and 10-fold cross validation, **Fig. S1E**) to predict the identity of the cells expressing 800 - 1500 genes (**Fig. S1F**). Only cells predicted as non-neuronal populations were included. This led to a dataset containing 26,429 neuronal cells and 26,913 non-neuronal cells.

### Clustering analysis of each major clusters

To further cluster the major clusters, the variable genes for each major cluster were identified with the “FindVariableGenes” function in Seurat. In this function, genes were placed into 20 non-overlapping bins according to their average expression and then a z-score between their dispersion (variance/mean expression) and the mean dispersion of the bin was calculated. The genes with the expression cpm > 1.5 and dispersion ≥ 1 standard deviation away from the expected dispersion in the corresponding bin were selected as variable genes. In some cases the threshold of dispersion was set to a higher value (up to 3 standard deviations) if the number of variable genes was larger than 2,500. Then the variable genes of each major cluster were used to calculate the top 30 principle components (PCs). Significant PCs with a p-value < 1e-3 were selected with JackStraw method in Seurat package. If too many PCs were significant, the top 10 PCs were selected. After principle component analysis (PCA), the coordinates of the cells in the significant PCs were used as input for cell clustering. For each major cluster, we first forced the “FindClusters” function to generate a large number of clusters by setting the resolution parameter to 3 or 4, then the marker genes of these clusters were calculated and any pair of clusters with less than 5 differently expressed genes (FC>2 and p-value < 0.01) were merged. Finally, we manually filtered out the cell clusters with substantial expression of 1) neuronal and non-neuronal marker or 2) established non-neuronal markers of different subtypes, since they potentially represent double droplets. We also excluded the cell clusters expressing high level of glutamatergic neuron markers (Slc17a6 and Slc17a7), since NAc does not contain glutamatergic neuron. In addition, cell clusters contributed by cells from less than three samples were also excluded. After these processes, our pruned dataset included 25,734 non-neuronal cells and 21,842 neuronal cells.

### Identification of cluster markers

The cluster-specific markers were identified by detecting the differentially expressed genes in the saline treated samples by comparing their expression in the given cluster versus the other clusters. Specifically, the cluster markers were calculated by using a likelihood ratio test with an underlying negative binomial distribution (*FindAllMarkers* function of the Seurat package) while controlling for the 1) number of UMI in each cell, 2) the percent of mitochondrial genes and 2) sample-to-sample variation.

### Cell clustering of scRNAseq data for MERFISH gene selection

Prior to selection of the MERFISH gene panel, the scRNAseq data was clustered in three successive rounds: 1) all cells classified into major cell types (neuronal and non-neuronal); 2) neuronal cells classified into major neuronal cell types (D1, D2, IN); and 3) each neuronal cell type classified separately into cell sub-types. For each round of clustering the raw counts for each cell were normalized to the total counts per cell and then logged (natural logarithm). Then, a group of highly variable genes was identified using the ‘highly_variable_genes’ function in Scanpy ^76^, an implementation of the z-scored normalized dispersion (variance/mean) method described previously ^77^, applying cutoffs for the minimum dispersion and minimum and maximum mean of 0.5, 0.025, and 4.0, respectively. Next, a simple linear regression was used to remove potential biases due to the total gene count or percentage of mitochondrial genes per cell, as implemented in the Scanpy ‘regress_out’ function. The data were then scaled to have mean of zero and unit variance. Then, a linear dimensionality reduction was performed using principal component analysis (PCA) on the identified highly variable genes and used the first 50, 44, 38, 28, and 26 principal components (PCs) for the all-cells, neuronal-cells, D1, D2, and IN clustering, respectively. To determine the number of PCs to keep, we calculated the largest eigenvalue after randomly shuffling the gene values of the cell-by-gene matrix, and then kept all the PCs that had an eigenvalue larger than the median of the largest eigen values across 10 such shuffling iterations ^33^. Finally, cells were embedded in a *k*-nearest neighbor graph in PC space using Scanpy ^76^ for a range of k sizes before performing Louvain community detection ^78^ on each, as implemented in the python Louvain-igraph package. To select an optimal *k* value, we used a bootstrapping analysis to evaluate cluster stability, as described previously ^33^, and selected a k value of 6, 30, 8, 8, and 8, for the all-cells, neuronal-cells, D1, D2, and IN clustering, respectively. Following the first round of clustering (all-cells), *Snap25* and *Gad1* were used to identify neuronal clusters to use in the second round of clustering. Following the second round of clustering (neurons), expression of *Ppp1r1b* and *Drd1* were used to identify D1 neurons, *Ppp1r1b* and *Drd2* to identify D2 neurons, and *Resp18* to identify interneurons (IN), which were each used in a third round of clustering to identify sub-types of each.

### Gene selection for MERFISH

To transcriptionally profile distinct cell populations with MERFISH, we designed a panel of 253 genes. Of these, 133 were chosen manually based on established markers for known cells types (excitatory and inhibitory neurons, oligodendrocytes, oligodendrocyte precursors, microglia, endothelial cells, ependymal, new-born neurons) or specifically pertinent to NAc biology (such as ion channels, ligands, receptors). To discriminate the sub-classes of the major neuronal cell types of the NAc (D1, D2, IN, see next section), we selected the remaining genes using a combination of two complimentary approaches: 1) mutual information (MI) analysis and 2) differentially expressed (DE) gene analysis. We selected the top 50 genes with most MI for each of the 3 groups of neuronal sub-clusters as described previously ^67^. Because of overlap between the three groups, this yielded a total of 114 genes. For DE genes, we included the top 2 DE genes for each sub-cluster when compared to the remaining cells in its respective group, as determined by the ‘rank_genes_group’ function of the Scanpy package, which employees a t-test to determine differential significance. This yielded 88 DE genes.

The combination of the genes selected manually and those selected by the data-driven approaches above resulted in a final gene panel of 253 genes. After screening the gene list to identify any genes that have a relatively high expression level, which are potentially challenging for MERFISH imaging ^33^, we selected 2 genes (Penk and Sst) to be imaged separately in a single two-color FISH imaging round, following the MERFISH run that imaged the remaining 251 genes.

### Design and construction of encoding probes

MERFISH encoding probes for the 251 genes were designed as described previously ^32, 33^. Each of the 251 genes was assigned to a unique binary barcode drawn from a 20-bit, Hamming Distance-4, Hamming-Weight-4 encoding scheme. We included 34 extra “blank” barcodes that were not assigned to any genes to provide a measure of the false-positive rate as described previously ^32, 33^. As previously described ^79^, we first identified all possible 30-mer targeting regions within each selected gene transcript. For each gene, we then randomly selected 60 of the 30-mer target sequences to comprise 60 encoding probes. For transcripts that were too short and had fewer than 60 targeting regions, we allowed the 30-mers to overlap by as much as 20 nt to increase the total number of probes targeting that transcript. We then assigned two readout sequences to each of the encoding probes associated with each gene. Each bit in the 20-bit code was associated with a unique readout sequence and for each gene, the readouts corresponding to the 4 “on-bits” of the gene’s assigned barcode, were evenly distributed over the entire set of its encoding probes. Finally, each encoding probe was flanked by the sequence of two PCR primers, the first comprising the T7 promoter and the second being a random 20-mer with homology to the encoding probes ^79^. Template DNA for encoding probes used for the 251 multiplexed genes was synthesized as a complex oligo pool (Twist Biosciences) and used to construct the final MERFISH probe set as described previously ^33^. Encoding probes for the 2 genes measured in sequential two-color FISH rounds were designed in a similar fashion as described above except: 1) 48 30-mer targeting sequences were chosen for each gene 2) one unique readout sequence was used for each gene, and 3) PCR primer sequences were omitted. Encoding probes were then synthesized in a 96-well plate format (Integrated DNA Technologies) and mixed to a suitable final concentration.

### Design and construction of readout probes

22 readout probes were designed to uniquely complement each of the 22 readout sequences used in the set of encoding probes. 20 of the 22 readouts correspond to the 20-bit barcode used for MERFISH imaging and the remaining 2 readout probes correspond to each of the 2 genes imaged in the sequential two-color FISH rounds. Each of the readout probes was conjugated to one of two dye molecules (Cy5, Alexa750) via a disulfide linkage, as described previously ^79^. Readout probes were synthesized and purified by Bio-synthesis, Inc., resuspended in Tris-EDTA (TE) buffer, pH8 (Thermo Fisher) to a concentration of 100 µM and stored at -20 ⁰C.

### Tissue slides preparation for MERFISH imaging

Wide type male C57BL/6N (8-10 weeks) mice were used for preparing striatal slices for MERFISH experiments. The animals were euthanized with isoflurane and the brains were quickly harvested and rinsed with ice-cold PBS. Each brain was cut into two hemispheres along the middle line and frozen immediately with dry ice in optimal cutting temperature compound (Tissue-Tek O.C.T.; VWR, 25608-930) and stored at -80 °C until slice cutting. The brains were cut into 10-µm-thick coronal sections with a cryostat (Leica CM3050S). Before slice cutting, the frozen brains embedded in OCT were put in the cryostat (-20 °C) for at least 30 min to stabilize the temperature. Slices were discarded until the NAc region was reached. Since there is no clear border between NAc and dorsal striatum, the brain tissues were trimmed to include both the NAc and dorsal striatum. To cover the entire NAc from anterior to posterior, a total of 14 10-µm-thick slices were taken at approximately 100 µm intervals.

The first 12 slices were distributed over 3 coverslips such that each of the 3 coverslips contained 4 slices spanning the anterior, mid, and posterior sections of the NAc (e.g. coverslip 1 received slices numbered: 1, 4, 7, 10, coverslip 2 received slices numbered: 2, 5, 8, 11). The fourth coverslip received slices numbered 13 and 14. Prior to slicing, the coverslips were prepared as described previously ^33^. Once placed on the coverslips, tissue slices were immediately fixed by incubating in 4% PFA in 1×PBS for 15 minutes at room temperature, which was followed by three successive washes with 1×PBS. Samples were then stored in 70% ethanol at 4 ⁰C for at least 18 hours to permeabilize the cell membranes. Under these conditions, the 4 coverslips could be stably stored for up to 2 weeks, with no appreciable degradation observed, while they were further processed in series for MERFISH imaging.

Next, the tissue slices were stained with the MERFISH probe set as described previously ^33^. Briefly, samples were washed three times with 2× saline sodium citrate (2×SSC, ThermoFisher AM9765), and equilibrated with encoding-probe wash buffer (30% formamide (ThermoFisher AM9342) in 2×SSC) for 5 minutes at room temperature. Wash buffer was then aspirated, and the coverslip was inverted onto a 50 µL droplet of an encoding-probe mixture on a parafilm coated petri dish. The encoding-probe mixture was comprised of ∼0.1 nM of each encoding probe used in the MERFISH rounds, ∼0.3 nM for each encoding probe used in the sequential two-color FISH round, and 1 µM of a polyA-anchor probe (IDT) in 2×SSC with 30% v/v formamide, 0.1% wt/v yeast tRNA (ThermoFisher AM7119) and 10% v/v dextran sulfate (Millipore S4030). Samples were stained for ∼48hr at 37 ⁰C. The polyA anchor probe contained a mixture of both DNA and locked nucleic acid (LNA) nucleotides (/5Acryd/ TTGAGTGGATGGAGTGTAAT T+TT+TT+TT+TT+TT+TT+TT+TT+TT+T, where “T+” denotes a thymidine LNA nucleotide and “/5Acryd/” denotes a 5’ acrydite modification) and was used to hybridize to the polyA sequence of polyadenylated mRNAs and anchor these RNAs to a polyacrylamide gel as described below. After staining, samples were incubated twice with encoding-probe wash buffer for 30 minutes at 47 ⁰C to remove excess and non-specifically bound probes. Samples were then cleared to remove background fluorescence as previously described ^33^. Briefly, samples were embedded in a thin 4% polyacrylamide gel and then incubated for 48 hours at 37 ⁰C in digestion buffer (50 mM Tris, pH 8.0 (ThermoFisher 15568025), 1mM EDTA (ThermoFisher 15575020), 2% SDS (ThermoFisher AM9823), 0.5% Triton X-100 (Sigma T9284), and 1:100 proteinase K (NEB P8107S)), refreshing the buffer once after the first 24 hours. Following digestion, samples were washed four times with 2×SSC and stored at 4 ⁰C in 2×SSC supplemented with 1:1000 murine RNase inhibitor (NEB M0314L) prior to imaging.

### MERFISH Imaging

The homemade imaging platform was described previously ^80^. Prior to imaging, the sample was stained with a hybridization mixture containing the readout probes associated with the first round of imaging, and an Alexa-488-conjugated readout probe complementary to the poly-A anchor probe. The hybridization mixture was comprised of readout probes at a concentration of 3 nM in hybridization buffer: 2×SSC, 10% (v/v) ethylene carbonate (Sigma E26258), and 0.1% Triton X100 (Sigma T9284). Staining incubated for 15 minutes at room temperature and sample was washed in hybridization buffer for 10 minutes at room temperature. Samples were then incubated with a solution of 10 µg/mL DAPI (ThermoFisher D1306) in 2×SSC for 5 minutes at room temperature, washed briefly in 2×SSC, and finally imaged.

For imaging, the sample coverslip was held inside a flow chamber (Bioptechs, FCS2) with a 0.75-mm-thick gasket (Bioptechs DIE# P47132), and buffer exchange within the chamber was directed using a custom-built automated fluidics system composed of three 12-port valves (IDEX, EZ1213-820-4) and a peristaltic pump (Gilson, MP3), configured as describe previously^32^. Imaging buffer was comprised of 2×SSC, 2 mM trolox (Sigma 238813), 50 µM trolox quinone, 5 mM protocatechuic acid (Sigma 37580), 1:500 recombinant protocatechuate oxidase 3,4-dioxygenases (rPCO 46852004, OYC Americas), 1:500 murine RNase inhibitor (NEB M0314L), and 5 mM NaOH (VWR 0583) to adjust final pH to 7.0. Cleavage buffer was comprised of 50mM TCEP (Gold Bio TCEP50) in 2×SSC.

Each imaging round consisted of readout probe hybridization (10min), washing with hybridization buffer (5min), flowing on imaging buffer (defined below), imaging of each field of view (FOV) (220 µm x 220 µm per FOV), readout fluorophore cleavage by cleavage buffer (15min), and washing with 2×SSC (5min). In the first round, we imaged the cell nucleus stained with DAPI and poly-A anchors stained with an Alexa488-labeled readout probe, at 7 focal planes separated by 1.5 µm in z. We then performed 12 rounds of two-color imaging, wherein the first 10 rounds imaged the barcoded-encoded RNA species (combinatorial smFISH rounds), the 11^th^ round imaged the individually labeled RNA species (sequential smFISH rounds), and the final 12^th^ round imaged the sample unlabeled to serve as measure of background fluorescence. For each combinatorial round, images were acquired with 750-nm and 650-nm illumination at 7 focal planes separated by 1.5 µm z to image the readout probes. For the sequential and background rounds, images were acquired with 750-nm and 650-nm illumination at a single focal plane 3.5 µm above the glass surface. In addition, every imaging round described above also included a single z-plane image with 546-nm illumination to image the fiducial beads on the glass surface for image registration. Finally, the number of FOVs imaged varied for each sample base on the size and number of tissues present on the coverslip.

### Image analysis and cell segmentation

All MERFISH image analysis was performed using the MERlin (https://github.com/emanuega/MERlin) python package, which in turn uses algorithms similar to what has been described previously ^33, 80^. First, for each FOV, the images from each imaging round were aligned using their respective fiducial bead image to correct for X-Y drift in the stage position relative to the first round. For MERFISH rounds, images stacks for each FOV were then high-pass filtered to remove background fluorescence, deconvolved using 10 rounds of Lucy Richardson deconvolution, and finally low-pass filtered to account for small movements in the centroid of RNA spots across the imaging rounds. Individual RNA molecules were then identified using pixel-based decoding, as described previously ^79^. Following decoding, adjacent pixels that were assigned the same barcode were aggregated as putative RNA molecules, and then the list of putative RNA molecules with an area greater than 1 pixel was filtered, as described previously ^80^, to enrich for correctly identified transcripts with a gross misidentification rat of 5%. Single pixel spots were excluded because they are disproportionally prone to be spurious barcodes generated by random fluorescent fluctuations. Next, we identified cell segmentation boundaries for each FOV using a seeded watershed approach as described previously ^33^. The DAPI images were used as the seeds and the polyA signals were used to identify segmentation boundaries. Following segmentation, individual RNA molecules were assigned to individual cells based on if they fell within the segmented boundaries. For the sequential two-color FISH rounds, images were high-pass filtered and the expression level of each gene in each cell was calculated as the sum of the fluorescence intensity of all pixels within the segmentation boundary of the central z plane of each cell. Finally, the signals from the 2 genes from the sequential two-color FISH round were merged with the RNA counts matrix from the 251 genes measured in the MERFISH run and used for cell clustering analysis.

### Cell clustering analysis of MERFISH data

The cell-by-gene matrix as obtained above was preprocessed with several steps. First, to remove spurious segmentation artifacts, we removed segmented “cells” that had a volume less than 100 µm^3^ or larger than 4000 µm^3^, which was chosen empirically to ensure retention of some larger inhibitory cell sub types that were observed in subsequent analysis. Because the expression units of the 251 genes measured in the MERFISH rounds was fundamentally different from that of the 2 genes measured in sequential FISH rounds (integer counts per cell versus total fluorescence intensity per cell, respectively), preprocessing of these two types of measurements required a different set of steps. For the counts matrix from the 251 genes the steps were as follows: 1) To reduce potential noise introduced by cells with few measured RNA molecules, we removed cells with fewer than 5 total RNA counts; 2) To normalize for variation in cell size, owing to either true biological differences between cell types or states, or to the fact that the entire soma of every cell did not occupy the 10-µm thick tissue slice, we divided the RNA counts per cell by the imaged volume of each cell; 3) To correct for minor batch fluctuations in measured RNA counts across slices, we normalized the total counts per cell to the same value (100 in this case). For the intensity matrix from the 2 sequential-round genes, we applied two steps to remove background fluorescence. 1) To equalize the baseline fluorescence for each gene across all slices, within each slice, the median intensity per gene was subtracted from each cell; 2) The final two-color imaging round was “blank” in that no readout probes were added, providing an autofluorescence background measurement for each cell in each imaging channel. This background was subtracted for each gene in each cell according to its imaging color channel.

After the preprocessing described above, we applied three successive rounds of clustering analysis to identify cell types and sub types. The first round was applied to all cells to identify major non-neuronal, neuronal and interneuron (IN) populaitons. In the second round we selected only the medium spiny neurons (MSNs) identified in the first round to further discriminate Drd1 positive (D1) and Drd2-positive (D2) neurons. In the first two rounds of clustering, established marker genes were used to identify cell populations: *Ppp1r1b* (MSN), *Resp18* (IN), *Snap25* (other-neuronal), *Aldoc* (astrocytes), *C1qa* (microglia), *Mobp* (oligodendrocytes), *Pdgfra* (oligodendrocyte precursors), *Rgs5* (endothelial), *Ccdc153* (ependymal), *Drd1* (D1 MSN), *Drd2* (D2 MSN). In the third round the D1, D2, and interneuron (IN, identified in the first round) populations were each clustered separately into the final reported sub-types. Each round of clustering followed the same basic workflow, similar to what was done for the clustering of the scRNA-seq data prior to MERFISH gene panel selection described above. Briefly, the cell-by gene expression matrix was log transformed and each gene was scaled to have unit variance and a mean of zero. Next, PCA was applied, using all genes, to reduce dimensionality and the number of PCs to keep were selected in each instance (26, 21, 22, 25, and 28 for all-cells, MSN, D1, D2, and IN clustering, respectively) as described above and previously ^33^. Then, as described above, cells were embedded in a *k*-nearest neighbor graph in PC space and clustered using Leiden community detection ^81^ as implemented in the Leidenalg python package. To remove an apparent batch effect between the datasets from each of the two mouse brains used, we employed the Harmony algorithm ^82^, as implemented in the Harmonypy python package, to correct PCs in the MSN, D1, D2, and IN analysis prior to clustering. Optimal *k* values of 25, 12, 10, 25, and 45 were chosen for the all-cells, MSN, D1, D2, and IN clustering, respectively, using the bootstrapping method described above. Finally, we manually filtered the clustering results by checking the gene expression and spatial distribution of each cell type (subtype), to remove cell clusters representing double-lets (based on co-expression of established makers of multiple cell types) or located out of striatum.

### Integration of scRNA-seq and MERFISH data

This analysis were done using the Seurat R package (v 3.2.0) ^83^ in combination with the R version of the Harmony algorithm ^82^ (v 1.0, https://github.com/immunogenomics/harmony) and some customized R scripts. We used the list of MERFISH genes to build a combined count matrix between the scRNA-seq counts and the normalized MERFISH RNA counts. Next, the counts were normalized to the library size, log-scaled and their batch un-corrected PCA was calculated using the Seurat functions: “NormalizeData”,“FindVariableFeatures”,“ScaleData” and “RunPCA” respectively. The “RunHarmony” function was then used with default parameters for batch correction in the PCA space. Next, The top 20 batch corrected PCs were used to generate the UMAP embeddings. The same approach was followed to respectively integrate D1 and D2 MSN data between scRNA-seq and MERFISH.

### Estimation of the similarity between scRNA-seq and MERFISH clusters

As MERFISH clusters captures both the transcriptional and spatial distribution, we used the MERFISH identified clusters as our reference. The degree of correspondence between scRNA seq and MERFISH clusters were calculated based on the k-nearest neighbors classification. Given the top 20 batch corrected PCs from Harmony, we identified for each scRNA-seq cell its 30 nearest MERFISH cells using the “Seurat:::FindNN” function with parameters “cells1 = scRNACells, cells2 = merFISHCells,, internal.neighbors = NULL, dims = 1:20, reduction = “harmony”, nn.reduction = “harmony”, k = 30, nn.method = “rann”, eps = 0”, then calling the function Seurat::GetIntegrationData to get the results. A scRNA-seq cell was a given MERFISH cluster label if more than 70% of its MERFISH neighbors belong to the same MERFISH cluster, otherwise it was considered as ambiguous. The degree of similarity between scRNA-seq and MERFISH cell clusters is then calculated as the proportion scRNA-seq cells that could be matched to each MERFISH cluster for each scRNA-seq cluster. From the total scRNA-seq data, only 3.04% of scRNA-seq cells could not be confidently matched to a MERFISH cluster, while 24.97% and 21.45% of scRNA-seq D1 and D2 MSN cells could not be confidently matched to a corresponding D1 or D2 MERFISH cluster.

### Prediction of the enrichment of GWAS genes in MERFISH clusters

To identify the MERFISH clusters that could specifically express GWAS genes not included in the 253 MERFISH gene panel, we first selected all the scRNA-seq cells that could be given a MERFISH labels. The enrichment of GWAS genes in the corresponding scRNA-seq data was performed as previously described ^84^. Briefly, from the NHGRI-EBI GWAS catalog (version 1.0.2) ^85^, we got the list of GWAS candidate genes and their associated disease. Only GWAS genes having an exonic mutation were considered. Next, we calculated the mean expression of each gene in each cluster excluding outlier values (expression < 99^th^ percentile). To annotate the enriched (1) or non-enriched (0) clusters for each GWAS gene, we ranked each gene for the mean expression in each cluster in an increasing order and the clusters having an average expression larger than knees value in the plot were marked as expressed. (kneepointDetection method, SamSPECTRAL package ^86^.

### Estimation of cluster density in a MERFISH slice

To visualize the spatial density distribution of a MERFISH cluster or group of clusters in a MERFISH slice we used the Gaussian kernel density estimator implemented in the function embedding density from the python module Scanpy (v1.6.0) ^87^.

### Generation of plots and heatmaps

R and Python were used interchangeably to generate the different plots and heatmaps. Heatmaps were generated using the R/Bioconductor package ComplexHeatmap ^88^ or the pheatmap R package ^89^. All the other plots were generated using the ggplot2 package ^90^. Dimensional embedding plots were mainly generated using Seurat R package (v 3.9.9.9008) ^83^. Scanpy (v1.6.0) ^87^ was also used to generate some gene expression heatmaps, dotplots and the density plots.

### Interactive visualization of MERFISH data

To enable the interactive exploration of MERFISH data, we built upon the open-source version of cellxgene v0.15.0 (https://github.com/chanzuckerberg/cellxgene). We did some additional customization to the color schemes and highlight behavior to fit our desired behavior. The actual version can be accessed at: http://35.184.4.122:5050/ (MERFISH) and http://35.184.4.122:9090/ (scRNA-seq).

### Data and code availability

The scRNA-seq data has been deposited to GEO with accession number: GSE118020. The MERFISH data is available at the Brain Image Library (https://download.brainimagelibrary.org/fc/4c/fc4c2570c3711952/). Code for MERFISH image acquisition is available at https://github.com/ZhuangLab. Code for MERFISH image analysis is available at https://github.com/ZhuangLab/MERlin. Code for MERFISH and scRNA-seq integration is available at https://github.com/YiZhang-lab/NAcMERFISHscRNAseqAnalysis.

## Acknowledgements

We thank Dr. Zhiyuan Chen and Mr. Qiangzong Yin for their help with sequencing; Dr. Bernardo L. Sabatini for critical reading of the manuscript. We thank the support of the HMS Neurobiology Imaging Facility (supported by NINDS P30 Core Center Grant #NS072030) and its staff Ryan Carelli and Michelle Ocana; Mouse Behavior Core of Harvard Medical School and its director Dr. Barbara Caldarone. This project was supported by the National Institutes of Health (NIDA 1R01DA042283 to Y.Z.; NIMH U19MH114821 to X.Z.), the Silicon Valley Community Foundation (to Y.Z.), and the Howard Hughes Medical Institute. Y.Z. and X.Z. are investigators of the Howard Hughes Medical Institute.

## Author Contributions

Y.Z. conceived the project; Y.Z. and X.Z. supervised the project; R.C, T.R.B., Y.Z. and X.Z. designed the experiments; R.C. performed the single-cell RNA-seq experiments; T.R.B. and J.H. performed the MERFISH experiments; M.N.D analyzed the single-cell RNA-seq data; T.R.B. and M.N.D. analyzed the MEFFISH data; A.B. and R.C. performed the RNAscope experiments; W.C. and L.M.T. helped with preparing scRNA-seq samples; R.C, T.R.B., Y.Z. and X.Z. interpreted the data; R.C., T.R.B., M.N.D, Y.Z. and X.Z. wrote the manuscript.

## Competing Interests

X.Z. is a co-founder and consultant of Vizgen.

## Extended data figure legends

**Extended Data Fig. 1.**
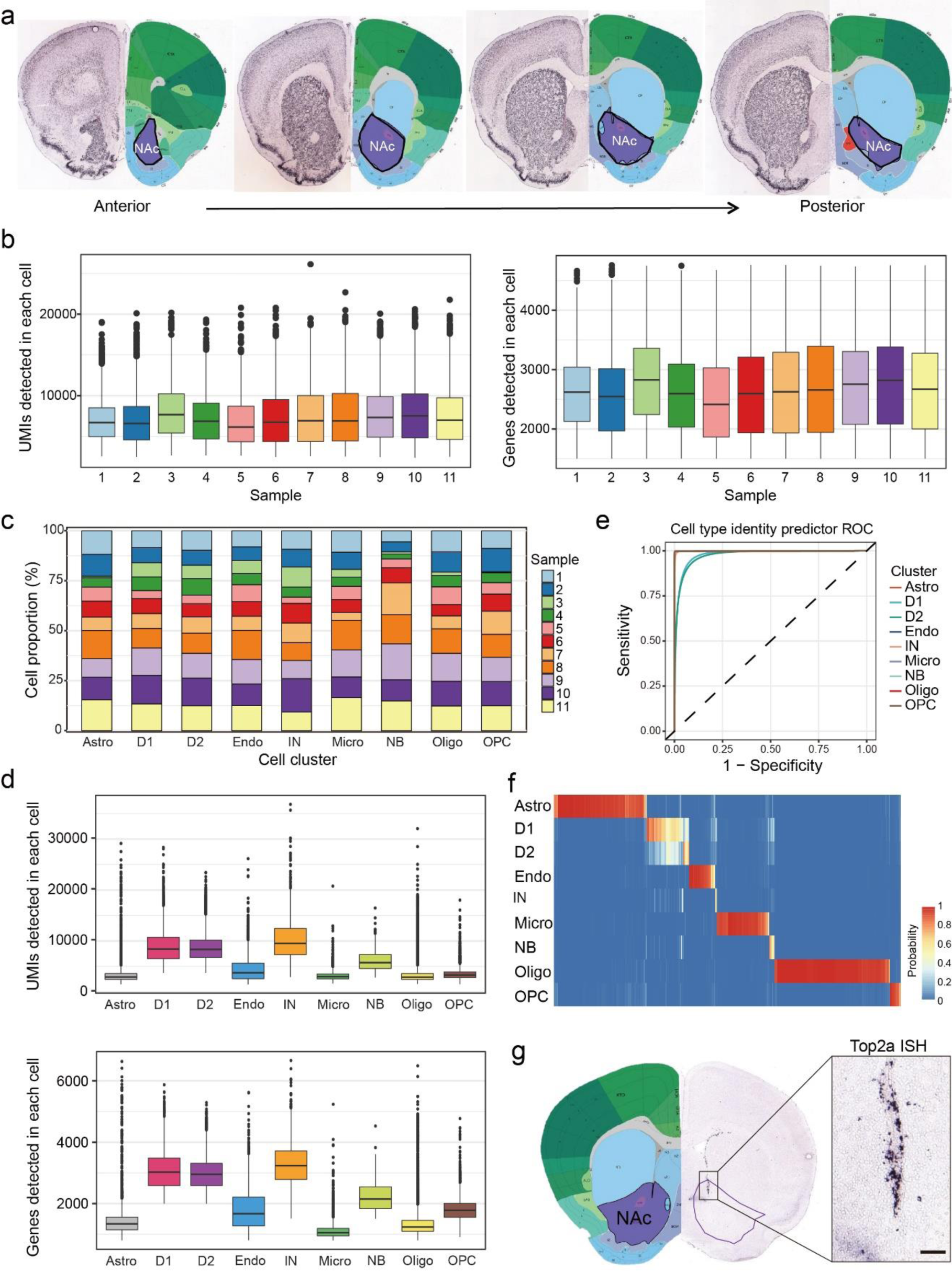
Identification of the transcriptionally distinct cell types in mouse NAc. **a,** Schematic diagram showing the nucleus accumbens region used for single-cell RNA sequencing. Adult mouse brain was first cut into 0.5mm-thick serial coronal sections and then NAc tissues (shown in purple contours) were dissected from successive slices along the rostral caudal axis. The brain pictures were taken from Allen Mouse Brain Atlas. **b,** Bargraphs showing the distribution of UMI and gene number detected in each cell across the 11 samples. Different samples are color-coded. **c,** Histograms showing the percentage of cells from each sample that contribute to the major cell clusters. Different samples are color-coded. **d,** Bargraphs showing the distribution of UMI and gene number detected in each cell across the 9 major cell clusters. Different cell clusters are color-coded. Astro, astrocyte; D1, medial spiny neuron, D1-receoptor subtype; D2, medial spiny neuron, D2-receptor subtype; Endo, endothelial cell; IN, interneuron; Micro, microglia; NB, neural stem cells and neuroblast; Oligo, oligodendrocyte; OPC, oligodendrocyte progenitor cell. **e,** ROC curves showing the high accuracy of the cell identity predictor, especially for non neuronal cells. The curves of different cell types are represented by different colors. **f,** Heatmap showing the results of predicted identity of cells with 800 to 1,500 genes detected. The prediction probability is color-coded. **g,** *In situ* hybridization of the cell-cycle gene, *Top2a*, showing the distribution of neural stem cells and neuroblasts in the ventral wall of the lateral ventricle. The purple contour indicates the NAc, and the boxed region in the left panel is enlarged and shown on the right. The ISH data was obtained from Allen Mouse Brain Atlas. Scale bar, 100 µm.

**Extended Data Fig. 2.**
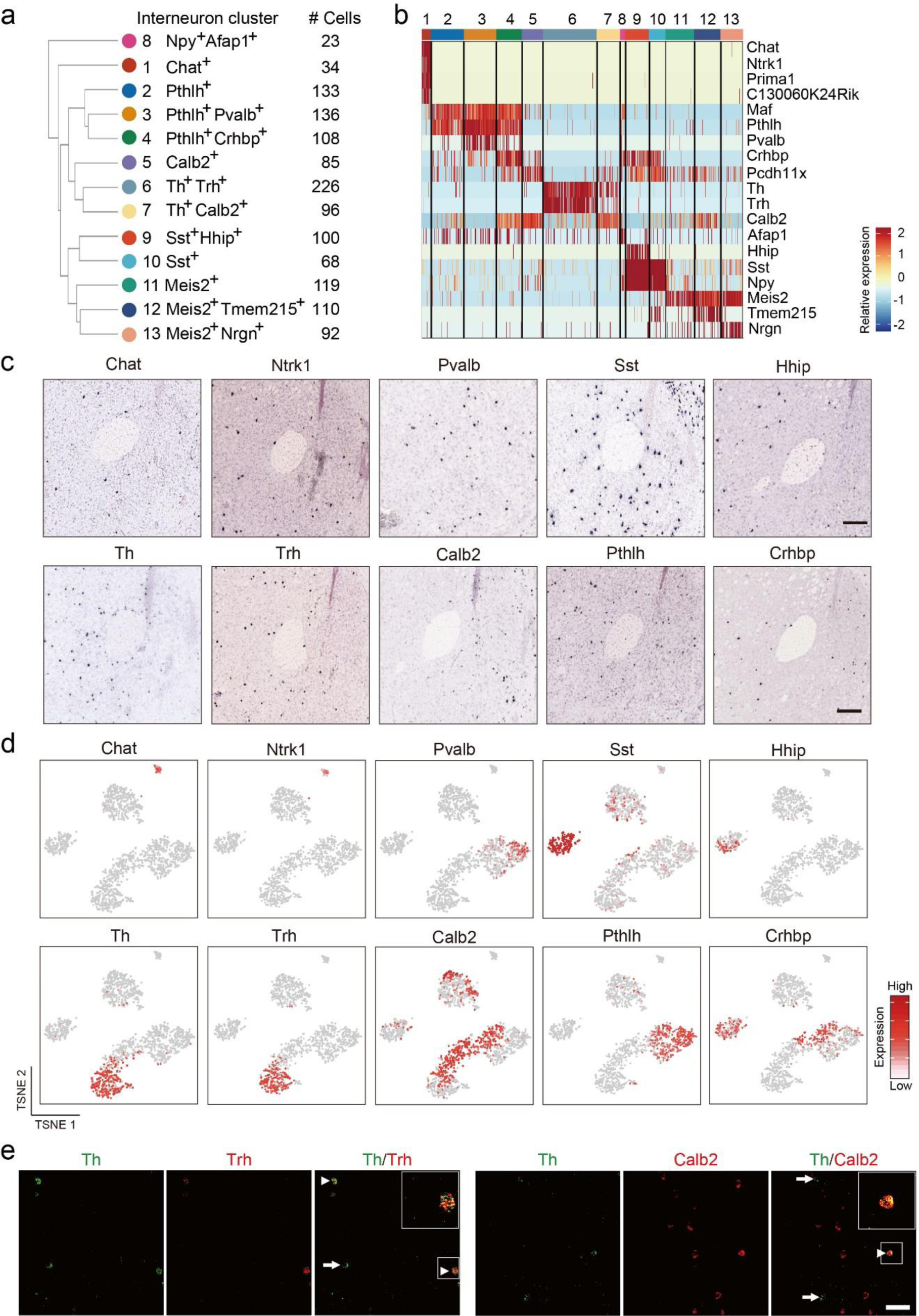
Gene expression and spatial pattern of NAc interneuron subtypes. **a,** Hierarchical relationship of the 13 NAc interneuron subtypes identified from scRNA-seq. The dendrogram indicate the relatedness among interneuron subtypes based on their gene expression. The markers and the number of cells of each subtypes are shown. **b,** Heatmap showing the expression pattern of interneuron markers across different subtypes. Each column represents a single cell. The gene expression level is color coded. The 13 interneuron subtypes are indicated by different colors on top. **c,** ISH showing the expression of selected interneuron subtype markers identified by scRNA-seq in NAc. The regions around the anterior commissure were shown except for Pvalb, which was more enriched in lateral part of NAc. The images were obtained from the Allen Mouse Brain Atlas. **d,** tSNE plots showing the expression of interneuron subtype markers across NAc interneuron subtypes. The gene expression level is color-coded. **e,** FISH showing the overlap of selected interneuron markers in NAc. Arrow heads indicate the cells co-express the two genes. Arrows indicate cells expressing one marker gene. Scale bar, 50 µm.

**Extended Data Fig. 3.**
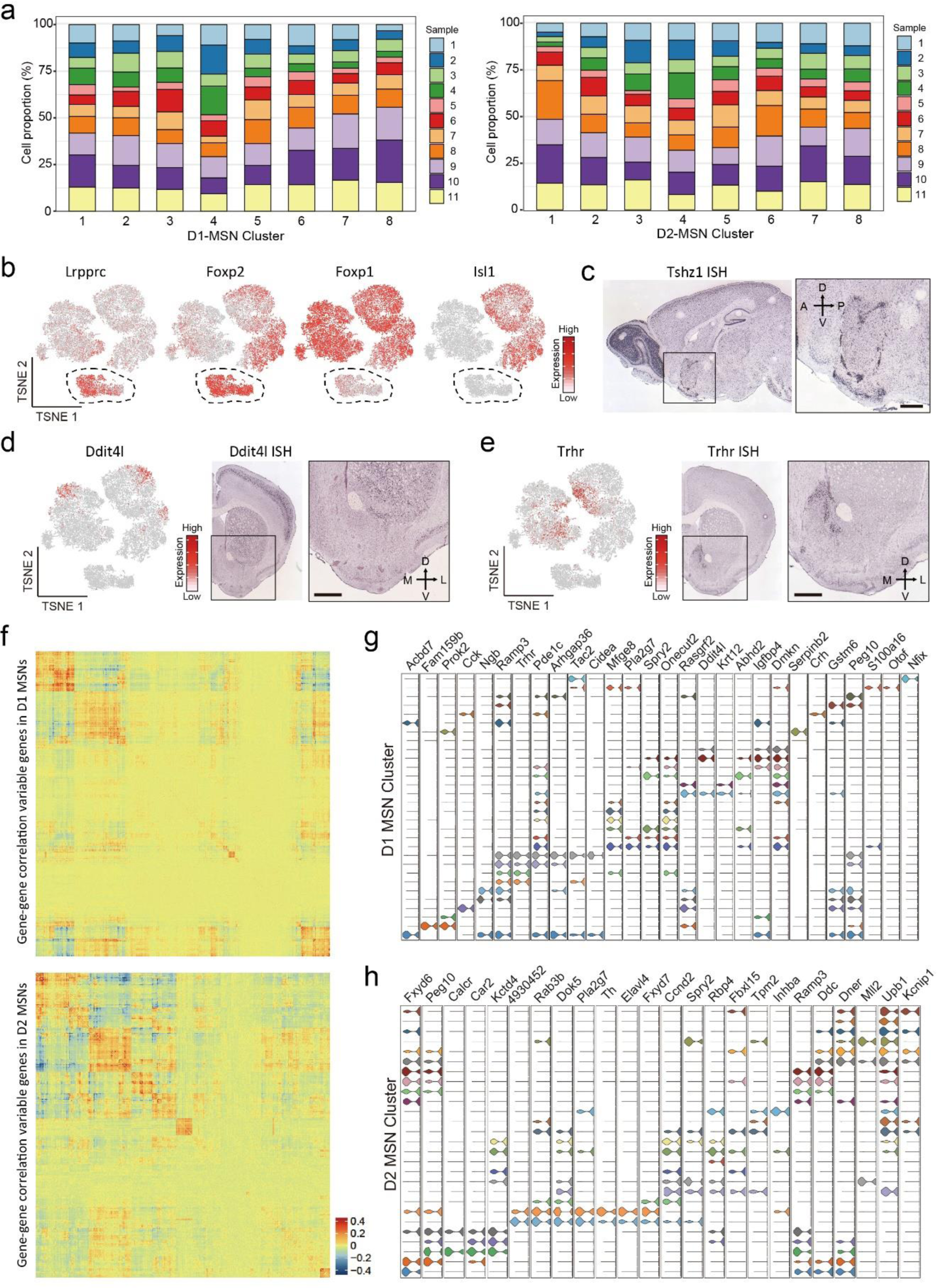
Molecularly defined MSN subtypes exhibit distinct spatial distribution in NAc. **a,** Histograms showing the percentage of cells from each of the 11 samples across D1 and D2 MSN subtypes. Different samples are represented in different colors. **b,** tSNE plots showing the expression pattern of *Lrpprc*, *Foxp2, Foxp1* and *Isl1* across MSN populations. The expression level is color-coded. **c,** ISH of *Tshz1* showing the distribution of *Tshz1*^+^ cells in NAc. Sagittal section of mouse brain including NAc is shown. The boxed region is enlarged and shown on the right. Data are obtained from Allen Mouse Brain Atlas. Scale bars, 500 µm. **d, e,** *Ddit4l* (d) and *Trhr* (e) are enriched in subpopulation of MSNs. Left panel, tSNE plots showing the expression pattern of *Ddit4l* (d) and *Trhr* (e) across MSN populations. The expression level is color-coded. Right panels, ISH image showing the distribution of *Ddit4l*^+^ (d) and *Trhr*^+^ (e) cells in NAc. The boxed regions are enlarged and shown on the right. Data is obtained from Allen Mouse Brain Atlas. Scale bar, 500 µm. **f,** Heatmaps showing the gene expression correlation of differentially expresses genes in D1 MSNs (upper panel) and D2 MSNs (lower panel). The correlation is color-coded, and genes were sorted into groups with higher intra-group correlation. **g, h,** Violin plots showing the expression of selected markers across high-resolution D1 (**g**) and D2 (**h**) MSN subtypes. Different MSN subtypes are color-coded. The mRNA level is presented on a log scale and adjusted for different genes.

**Extended Data Fig. 4.**
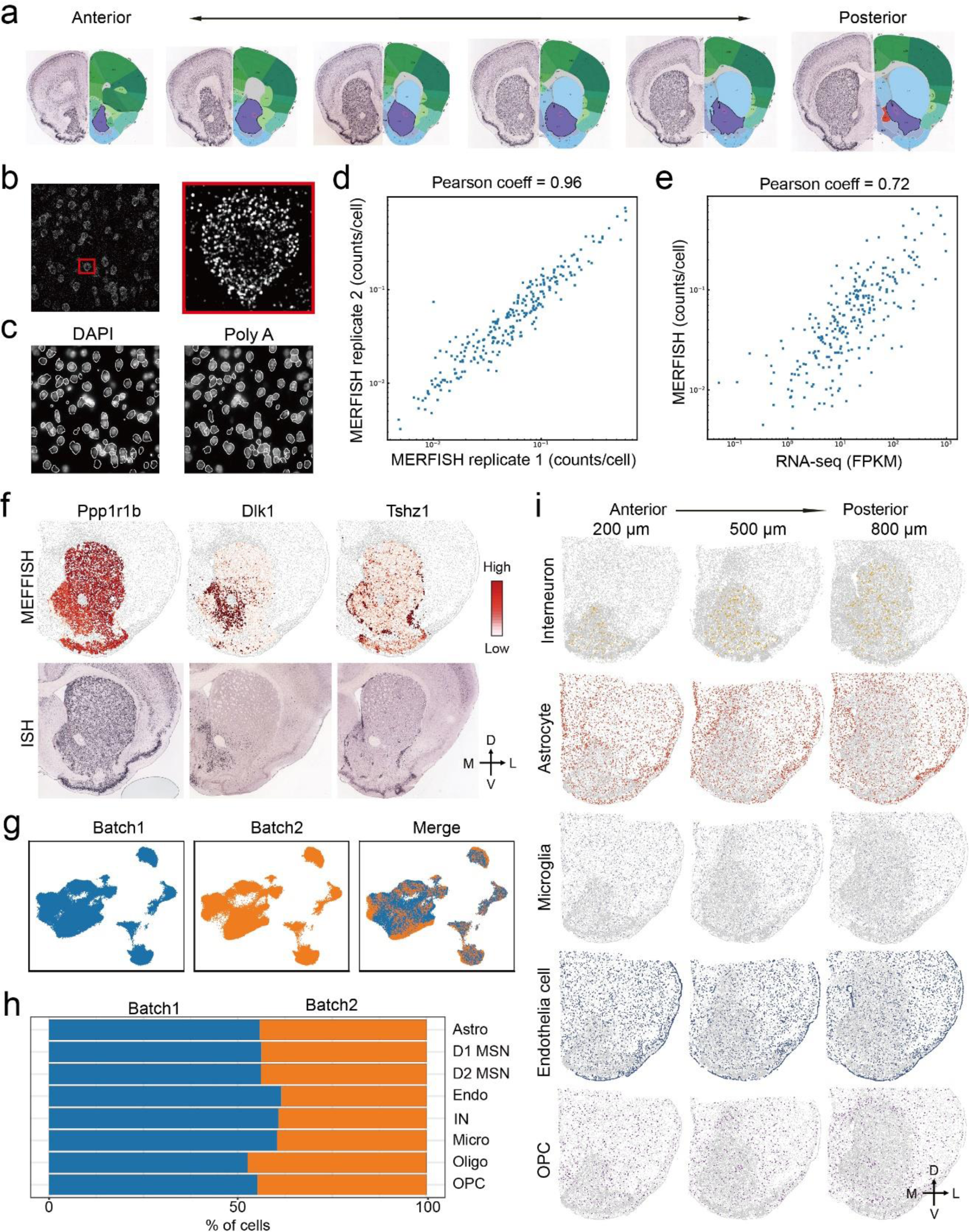
MERFISH revealed major cell types in striatum. **a,** Schematic diagram showing the striatal slices used for MERFISH. Adult mouse brain was first cut into 10 µm -thick serial coronal sections. Twelve brain slices with 100 μm interval between adjacent slices were used for MERFISH experiment. The brain pictures were taken from Allen Mouse Brain Atlas, note that only 6 slices were shown. **b,** One example image showing the maximum projection of images taken in one representative field-of-view (FOV) during MERFISH. The boxed region was enlarged and shown on the right. Individual RNA molecules were detected as single dots. **c,** DAPI (left) and poly(A) RNA (right) images were used to define the boundaries of each cell in white. The mRNA molecules detected were further assigned to different cells based on the cell boundaries. **d,** Scatterplot showing the average counts of each genes per cell detected by MERFISH in the two biological replicates. **e,** Scatterplot showing the average copy number of each genes per cell detected by MERFISH and bulk RNA-seq. The Pearson correlation coefficient is 0.77. **f,** Heatmap showing the expression pattern of selected genes in striatum as determined by MERFISH (upper panels), which are highly similar to the patterns determined by conventional ISH (lower panels). **g,** the Harmony algorithm ^82^ based UMAP plots showing the cells from the two replicates of MERFISH experiments with good overlap. **h,** Bar graph showing the proportion of major cell types from the two batches of MERFISH experiments after the Harmony integration. **i,** Spatial pattern of interneuron, astrocyte, microglia, endothelial cell and OPC in coronal brain sections at different anterior-posterior positions. Three of the twelve slices from a male mouse were shown. Colored dots were cells belong to the specified cell populations, while gray dots indicate all other cells. The 200, 500 and 800 μm labels indicate the distance from the anterior position (Bregma 1.94mm). The dorsal-ventral (DV) and medial-lateral (ML) axes are indicated.

**Extended Data Fig. 5.**
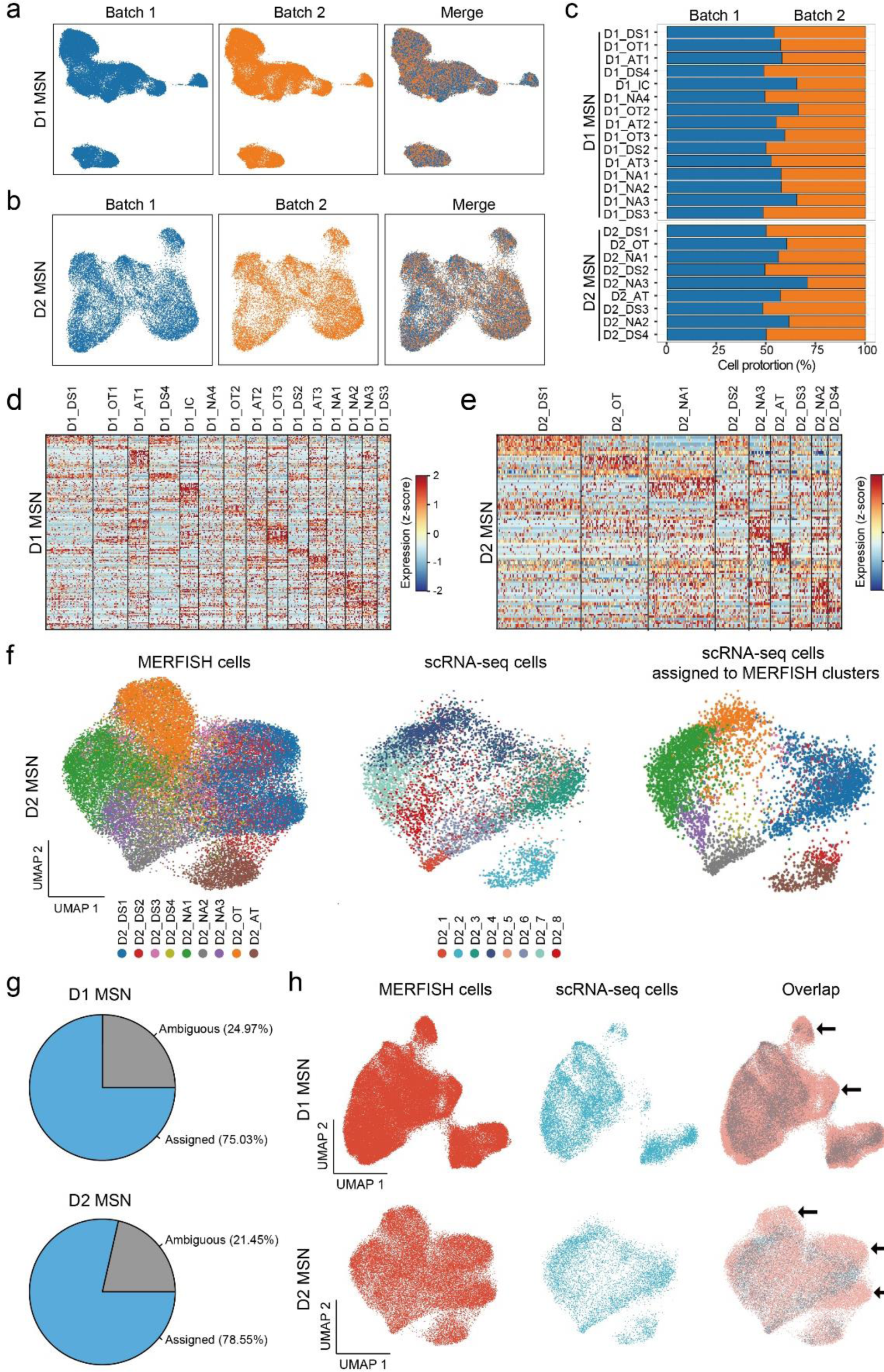
MERFISH identifies molecularly distinct D1 and D2 MSN subtypes in striatum. **a, b,** The Harmony algorithm ^82^ based UMAP plots showing the D1 MSNs (a) and D2 MSNs (b) from the two replicates of MERFISH experiments. Cells from the two batches occupy similar UMAP space. **c,** Bar graph showing the proportion of different D1 and D2 MSN subtypes from the two replicates of MERFISH experiments. **d, e,** Heatmaps showing the expression pattern of differentially expressed genes detected by MERFISH across D1 (**d**) and D2 (**e**) MSN subtypes. The expression level is color coded. The width of each column represents the abundance of each MSN subtype. **f,** Integrative analysis (Using the Harmony algorithm) of D2 MSNs from MERFISH and scRNA seq experiments. The D2 MSNs from MERFISH and scRNA-seq experiments were integrated into the same UMAP space. The initial identity of each cell was color coded and shown in the left and middle panels. Based on the nearest neighbors from the MERFISH experiments, the cells from scRNA-seq were assigned to one of the MERFISH D2 MSN subtypes shown on the right panel. **g,** Pie charts showing the percentage of D1 (upper) and D2 (lower) MSNs from scRNA-seq experiments which could or could not be assigned to a certain MEFFISH identify due to insufficient MERFISH k-NN belonging to the same cluster. **h,** The Harmony algorithm based UMAP showing the D1 MSNs (upper panels) and D2 MSNs (lower panels) from scRNA-seq and MERFISH experiments. The cells from different experiments were integrated into the same UMAP spaces. The arrows indicated UMAP spaces that are mainly occupied by cells from non-NAc region, thus the scRNA-seq cells were depleted comparing to MERFISH cells.

**Extended Data Fig. 6.**
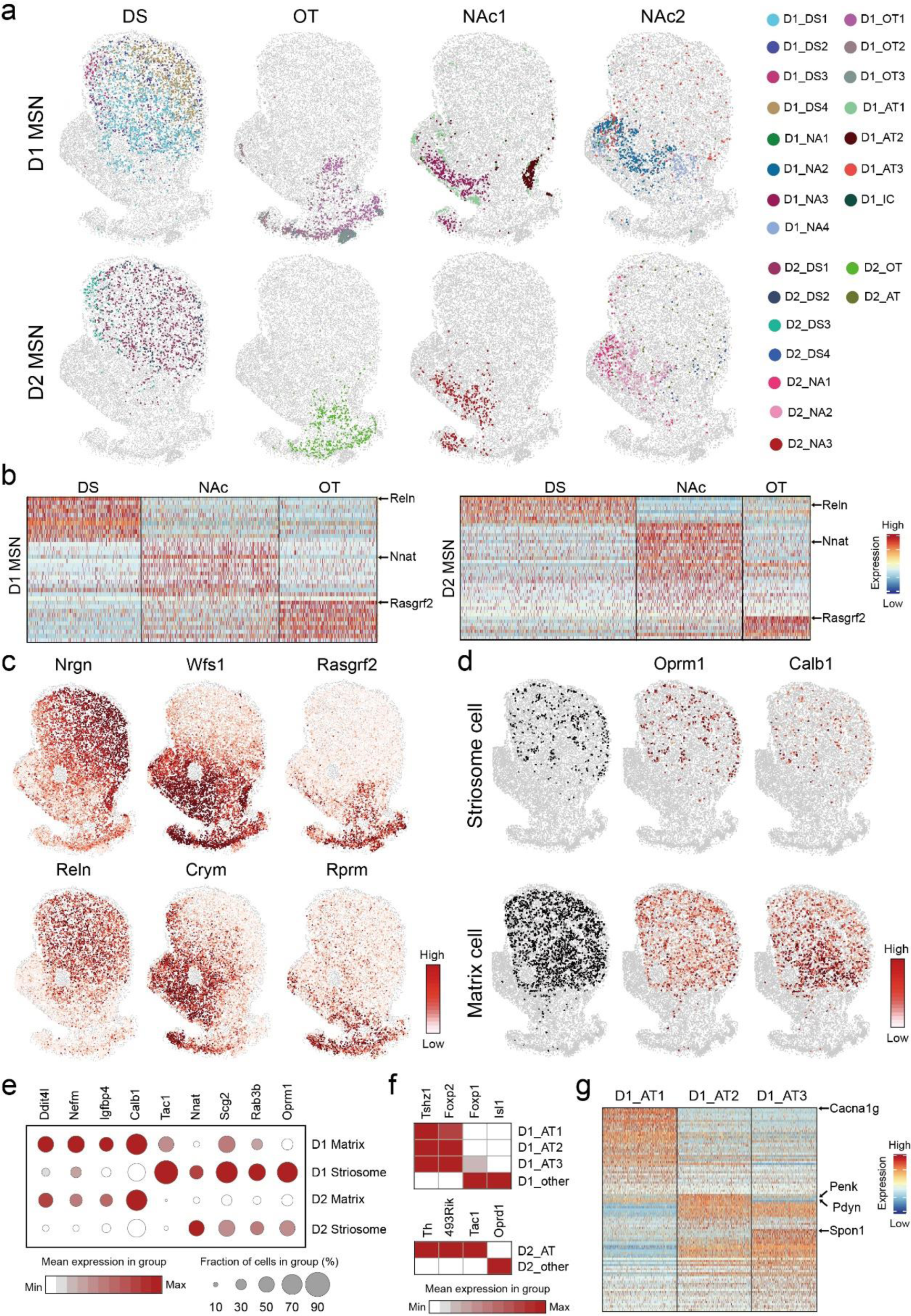
Molecular and spatial features of MSN subtypes underlie anatomic organization of striatum. **a,** Spatial patterns of different MSNs groups in striatal region. The MSN subtype groups are the same as Figure 6E. Different MSN subtypes are color coded. The D1 and D2 MSNs are shown in upper and lower panels, respectively. **b,** Heatmap showing the patterns of differentially expressed genes among D1 (left panel) and D2 (right panel) MSN subtype groups located in major striatal divisions. The MSN groups corresponding to different anatomic regions are the same as Figure 6E, but the two NAc groups are combined. *Reln, Nnat* and *Rasgrf2* are enriched in dorsal striatum, NAc and OT, respectively, and the patterns are shared in D1 and D2 MSN subtypes. **c,** The spatial heatmaps showing the expression pattern of *Nrgn, Reln, Wfs1, Crym, Rasgrf2, Rprm* in coronal sections, which are enriched in different striatal divisions. The expression level is color coded. **d,** Spatial and gene expression features of D1 MSN subtypes representing striosome (D1_DS2) and matrix (D1_DS1, D1_DS3 and D1_DS4) structure in dorsal striatum. The upper and lower panels show D1 subtypes representing striosome and matrix, respectively. The left panels show spatial pattern of D1 subtypes corresponding to striosome and matrix. The middle and right panels are heatmaps showing the expression of striosome enriched gene *Oprm1* and matrix enriched gene *Calb1* in these D1 MSN subtypes. **e,** Dotplot showing the expression pattern of selected genes in D1 and D2 MSN subtypes representing striosome and matrix. D1 Matrix: D1_DS1, D1_DS3 and D1_DS4; D1 Striosome: D1_DS4; D2 Matrix: D2_DS1 and D2_DS3; D2 Striosome: D2_DS2. The expression level is color coded. Dot size represents the fraction of cells in the subtype. **f,** Heatmap showing the expression of selected marker genes that distinguish atypical D1 and D2 MSN subtypes from other D1 and D2 MSN subtypes. The expression level is color coded. **g,** Heatmap showing the pattern of differentially expressed genes among the three atypical D1 MSN subtypes. Selected marker genes enriched in different atypical D1 MSN subtypes are labeled.

**Extended Data Fig. 7.**
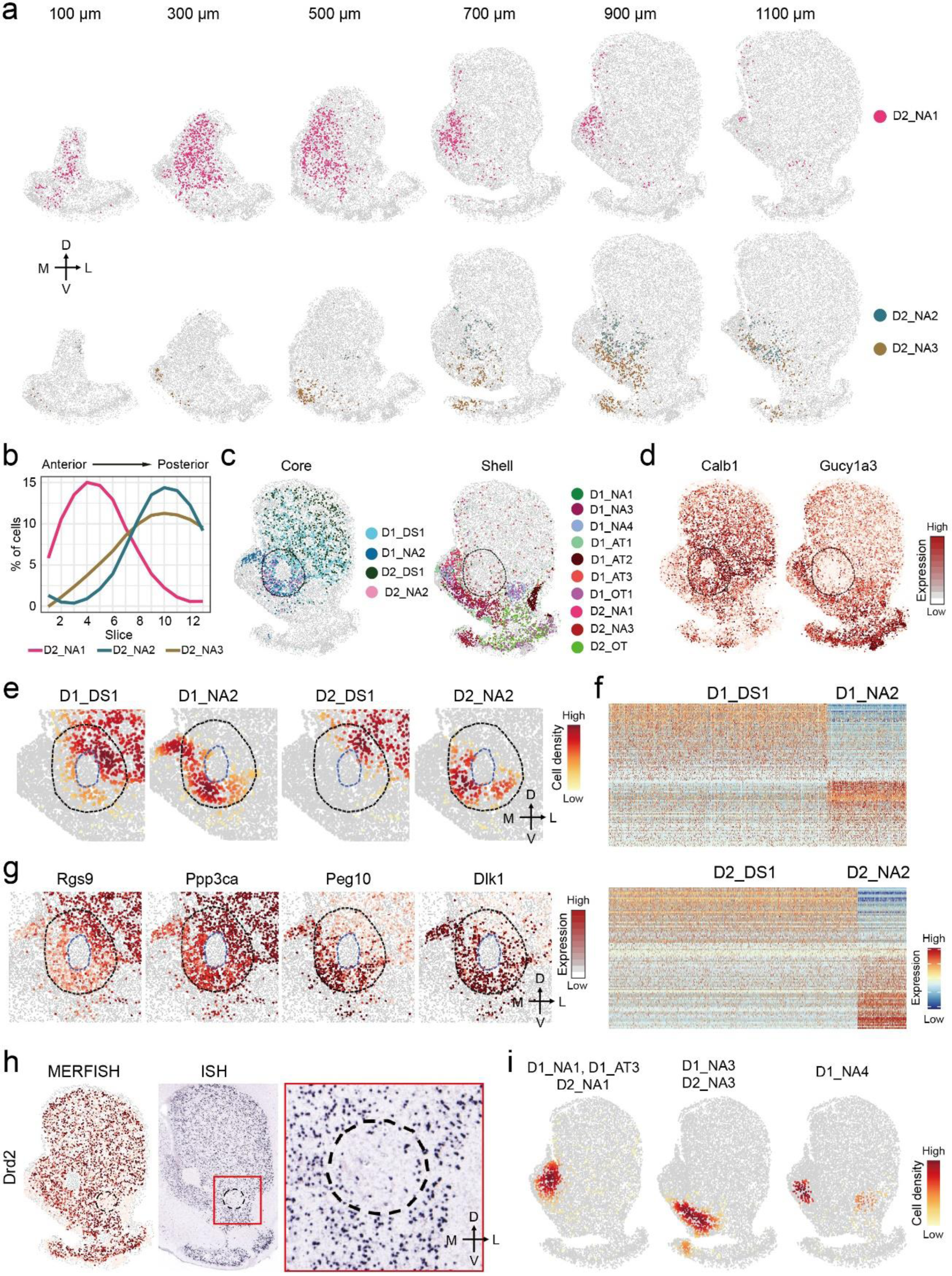
Molecular and spatial features of MSN subtypes underlie anatomic heterogeneity of NAc. **a,** Spatial patterns of selected D2 MSN subtypes in coronal sections at different anterior posterior positions. The subtypes enriched in anterior and posterior NAc are shown in upper and lower panels, respectively. Different subtypes are represented by different colors. The 100, 300, 500, 700, 900 and 1100 μm labels indicate the distance from the anterior position (Bregma 1.94mm). The dorsal-ventral (DV) and medial-lateral (ML) axes are indicated. **b,** The distribution of the three D2 MSN subtypes shown in (A) along the AP axis. **c,** The spatial pattern of MSN subtypes enriched in NAc core (left panel) or shell (right panel). Different subtypes are indicated with different colors. The core region is indicated by dashed line. **d,** Heatmap showing the expression of core enriched gene *Calb1* and shell enriched genes *Gucy1a3* in coronal brain sections. The expression level was color coded and the core region was indicated with dashed line. **e,** Density maps showing the enrichment of different D1 and D2 MSN subtypes in the dorsolateral and ventromedial part of NAc core. The same region as Figure 7F are shown. The NAc core and AC structure are labeled with dashed lines. The dorsal-ventral (DV) and medial lateral (ML) axes are indicated. **f,** Heatmaps showing the differentially expressed genes between D1 (upper panel) and D2 (lower panel) MSN subtypes enriched in different subregions of NAc core. The expression level is color coded. **g,** Heatmap showing the differential expression of *Rgs9, Ppp3ca, Peg10* and *Dlk1* between the dorsolateral and ventromedial part of NAc core. The same region as Figure 7F is presented. The NAc core and AC structure are labeled with dashed lines. The gene expression level is color coded. **h,** MERFISH and ISH detection of *Drd2* expression in striatum. The dorsal part of NAc lateral shell with low *Drd2* expression is labeled with dashed line. The boxed region in the ISH image is enlarged and shown on the right. The ISH data is from Allen Brain Atlas. **i,** Density maps showing the enrichment of different groups of MSN subtypes in distinct subregions in NAc medial shell.

**Extended Data Fig. 8.**
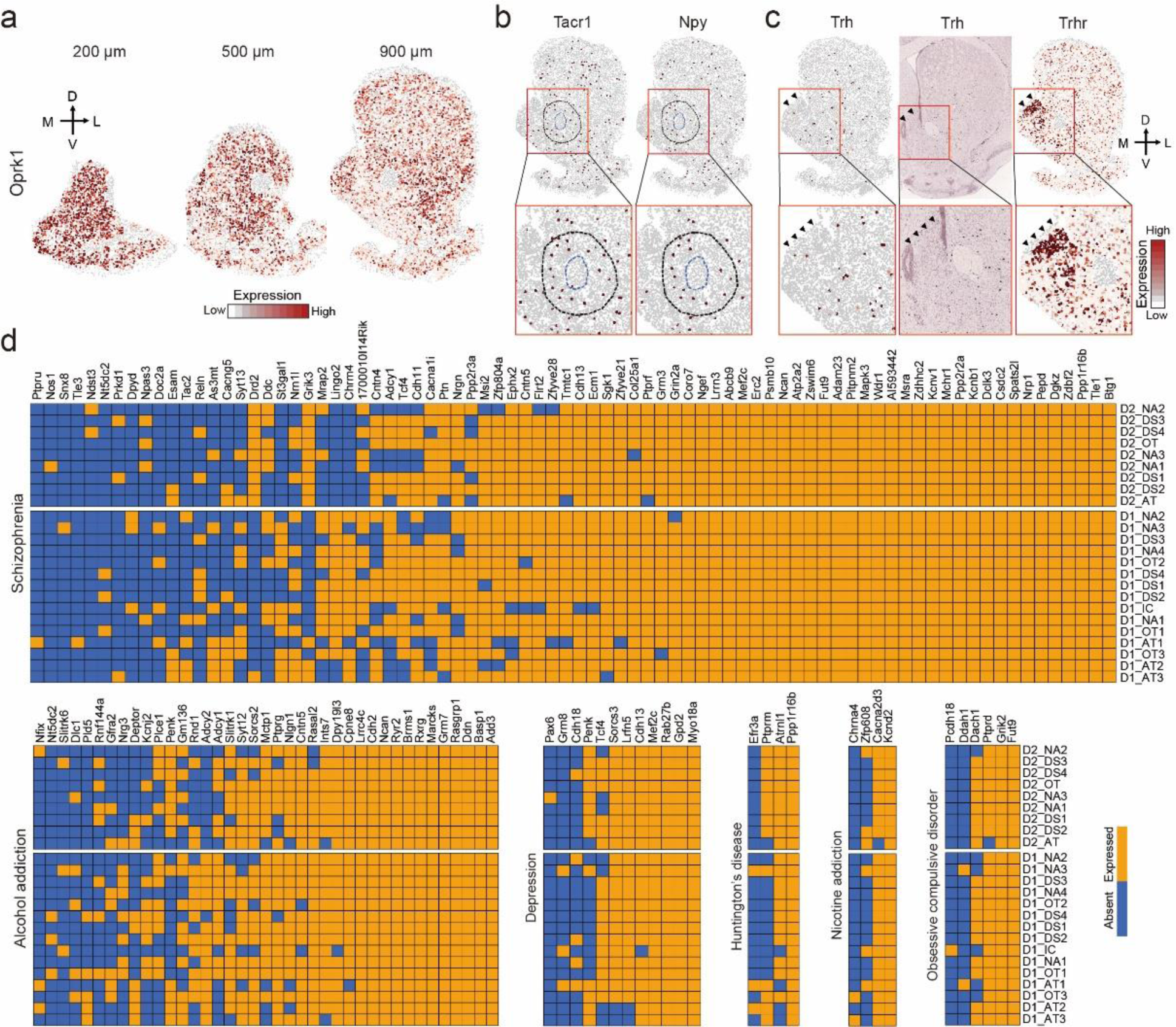
The molecular and spatial features of different neuron subtypes underlie their functional complexity. **a,** Heatmap showing the expression of *Oprk1* in coronal sections at different anterior-posterior positions. Note the *Oprk1* level is low in ventromedial part of posterior NAc. The expression level is color-coded. The 200, 500, and 900 µm labels indicate the distance from the anterior position (Bregma 1.94mm). The dorsal-ventral (DV) and medial-lateral (ML) axes are indicated. **b,** Heatmap showing the expression of *Tacr1* in *Chat*^+^ and *Npy*^+^ interneurons (left) and *Npy* in *Npy*^+^ interneurons (right). The boxed regions in the upper panels are enlarged and shown in lower panels. The NAc core and AC structures are labeled with dashed lines. **c,** Heatmap and ISH showing the expression pattern of *Trh* and *Trhr* in striatum. Arrowheads indicate the dorsoventral NAc with enriched *Trhr* expression. The boxed regions in the upper panels are enlarged and shown in lower panels. **d,** Heatmap showing the expression of GWAS candidate disease-relevant genes across different D1 and D2 MSN subtypes. Blue color, no enrichment; Orange color, gene enrichment. Genes associated with schizophrenia, alcohol addiction, depression, Huntington’s disease, nicotine addiction and obsessive compulsive disorder are shown.

